# DCX enables branching of subpellicular microtubules in *Plasmodium falciparum* gametocytes and is required for mosquito colonisation

**DOI:** 10.64898/2026.07.10.737674

**Authors:** Emma Ganga, Rosie Bridgwater, Amke Hackmann, Souradip Mukherjee, Olivier Mercey, Fergus Tollervey, Mufuliat Toyin Famodimu, Michael Delves, Paul Guichard, Virginie Hamel, Mathieu Brochet

## Abstract

*Plasmodium falciparum*, the causative agent of malaria, relies on specialised tubulin-based cytoskeletal structures to support its parasitic lifestyle. These include the conoid required for parasite motility and host-cell invasion, as well as subpellicular microtubules (SPMTs) that support parasite shape and rigidity. Here, we investigate the function of the doublecortin-domain protein DCX, a microtubule-binding protein previously associated with the *Plasmodium* conoid. We first show that, in *P. falciparum*, DCX is not expressed in the merozoite stage and is not required for the invasion of human erythrocytes. By contrast, DCX is expressed in ookinetes, the motile stage responsible for infecting the mosquito vector, where it associates with conoid tubulin fibres, consistent with a role in stabilising the conoid architecture. Unexpectedly, we find that DCX is required for *P. falciparum* transmission to the mosquito independently of conoid function. We further link this requirement to the distinctive organisation of SPMTs in *P. falciparum* gametocytes, which display an unusual branching architecture comprising multiple microtubules of 15 to 18 protofilaments. Deletion of *DCX* leads to a reduction in SPMT branching and is associated with higher protofilament numbers, revealing a previously unrecognised role for DCX in shaping the ultrastructure of SPMTs in *P. falciparum* gametocytes. Altogether, our findings uncover the repurposing of DCX across distinct microtubule systems in transmission stages and identify DCX as a key factor mediating microtubule branching and stabilisation in SPMTs required for efficient mosquito transmission.

## Introduction

Malaria is caused by unicellular parasites of the genus *Plasmodium*, which belong to the phylum Apicomplexa. Their lifecycle spans both human and mosquito hosts and involves a wide range of cellular forms. These include intracellular replicative stages, extracellular motile or invasive forms, and quiescent intracellular stages, each characterised by distinct microtubule-based cytoskeletal architectures^1,2^. Microtubules are essential for a wide array of eukaryotic cellular processes, including cell division, cell motility, intracellular organisation, and organelle trafficking. They are polarised cylindrical polymers composed of protofilaments made of α- and β-tubulin heterodimers. Despite this conserved molecular scaffold, microtubules assemble into structurally and functionally distinct arrays, tailored to specific cellular contexts. Microtubules are classically described as 13-protofilament structures, a configuration most frequently observed in model organisms. However, a broad spectrum of microtubule architectures has long been recognised across eukaryotes^3^. Notable examples include the specialised microtubule arrays forming the suction disc of the intestinal parasite *Giardia lamblia* and the unusually large 40-protofilament microtubule found in mantidfly sperm^4^. Despite this diversity, our current understanding of microtubule biology remains heavily skewed toward metazoan systems.

Phylogenetically distant from traditional model organisms, *Plasmodium* displays a remarkable diversity of stable microtubule arrays, some conserved across eukaryotes, others uniquely tailored to its parasitic lifestyle. Like many eukaryotes, *Plasmodium* assembles axonemal microtubules that drive the flagellar motility of microgametes^5^, enabling them to reach and fertilise macrogametes, an essential step for initiating mosquito colonisation. In contrast, other microtubule structures are tailored to the parasitic lifestyle. For example, arrays of subpellicular microtubules (SPMTs) provide structural support and rigidity to invasive or motile forms of the parasite, such as merozoites, which invade erythrocytes; ookinetes, which traverse the mosquito midgut epithelium; and sporozoites, which are responsible for transmission to humans. Non-motile intracellular gametocytes that initiate mosquito transmission upon a blood feed also express SPMTs that confer their falciform pointed shape. Another distinctive cytoskeletal feature includes the tubulin fibres of the conoid complex^4^, a specialised apical structure that acts as a scaffold for the invasion machinery in motile stages^6^.

These structures display remarkable microtubule organisational diversity across developmental stages. The microgamete axoneme adopts the classical architecture of motile cilia, consisting of nine outer doublet microtubules surrounding a central pair of singlets^7^. Although this arrangement has not been formally characterised in *Plasmodium*, doublets typically comprise an incomplete 10-protofilament B-tubule attached to the side of a complete 13-protofilament A-tubule. Recent studies have shown that in *Plasmodium*, the assembly of the B-tubule within the basal body depends on δ-Tubulin^8^ and its attachment to the A-tubule requires at least two microtubule-associated proteins (MAPs), FAP52 and FAP20^9^.

In extracellular motile forms, SPMTs display a canonical organisation with 13 protofilaments. However, these microtubules show different profiles depending on their decoration by MAPs^4^. For example, sporozoite and ookinete SPMTs show Interrupted Luminal Helices (ILH) made of an array of proteins, including TrxL1 and SPM1, which may explain their flattened cross-section compared with the circular section of merozoite SPMTs that lack these proteins^4^. SPMTs of *P. falciparum* gametocytes show a unique organisation with 13 to 18 protofilament fibres that form rafts of up to five branched microtubules^4^. While all zoite SPMTs have the same polarity with minus ends at the apical end, gametocyte SPMTs show an apparent random polarity, suggesting assembly from both ends of the cell. These morphological features of gametocytes are not conserved across the *Plasmodium* lineage, with most gametocytes from rodent, bird, and primate parasites being round and not expressing SPMTs^10^. This diversity in protofilament numbers and branched organisation of *P. falciparum* gametocyte SPMTs is unprecedented, and the corresponding biogenesis and physiological relevance remain to be elucidated.

Finally, the tubulin fibres of the conoid complex also show a unique organisation. While *Plasmodium* was initially thought to lack a conoid, we recently discovered that the ookinete stage expresses a reduced tubulin-based conoid^11^. This was confirmed by *in situ* Cryo-ET, which revealed that the ookinete conoid is made of open comma-shaped tubulin fibres^4^. In *Toxoplasma gondii*, a widely studied apicomplexan parasite responsible for toxoplasmosis in humans, the conoid has been most thoroughly characterised. In this parasite, conoid fibres are initially assembled as canonical 13-protofilament microtubules and undergo a maturation process that transitions them into a distinctive nine-protofilament left-handed comma-shaped structure^12^. This atypical shape and stability of conoid fibres have been attributed to the DCX protein, also referred to as Apicortin^13,14^, which is conserved across apicomplexan parasites. DCX shows a doublecortin domain and a TPPP/P25-α domain, both of which are known modulators of tubulin polymer structure. Consistent with this role, heterologous expression of *T. gondii* DCX in *Xenopus* cells promoted the formation and stabilisation of curved microtubule fibres^15^. A recent atomic model of the *Toxoplasma* conoid showed that DCX forms a meshwork over the crests of protofilaments 5–9 with conoid protein hub 1 (CPH1) and CF6^16^ where DCX binds to the tubulin lattice in a manner distinct from its mammalian orthologs^17^. In *Plasmodium*, the role of DCX remains less clear. While earlier studies suggested a function for DCX in *P. falciparum* asexual blood-stage replication, more recent work demonstrated that, despite its expression at the apex of ookinetes and sporozoites, DCX was dispensable throughout the life cycle of *P. berghei*, a parasite infecting rodents^14^.

In this study, we first show that DCX is neither expressed nor required for the proliferation of *P. falciparum* asexual blood stages. Unexpectedly, however, DCX proves important for the successful transmission of *P. falciparum* to the *Anopheles* mosquito vector as opposed to its *P. berghei* orthologue. Using iterative ultrastructure expansion microscopy (iU-ExM), we confirm its association with conoid fibres in ookinetes, but attribute its primary essential function to an earlier, conoid-independent role during sexual development. Specifically, we detect DCX expression in early-stage gametocytes and link DCX to the stabilisation of branched SPMT in stage IV gametocytes as well as the maintenance of axonemal symmetry and bundling in microgametes. Furthermore, *in situ* cryo-ET reveals that *DCX* deletion is associated with higher numbers of protofilaments in SPMTs, likely by stabilising open microtubules. Together, these findings reveal the multifaceted role of DCX in *P. falciparum* transmission, extending beyond its previously proposed function in conoid stability.

## Results

### *P. falciparum* DCX is neither expressed in merozoites nor required for the growth of asexual blood-stages

To investigate the role and localisation of DCX (PF3D7_0517800) in *P. falciparum*, we opted to conditionally delete the endogenously tagged *PfDCX* gene using the rapamycin-inducible DiCre system^18^. Briefly, a C-terminal 6xHA tag and two *loxP* sites, one within an exogenous 2*loxP*int intron and the other downstream of the 6xHA tag, were introduced in the endogenous *DCX* locus via double homologous recombination. These modifications enable conditional truncation of 256 out of 279 codons in the *PfDCX* gene upon rapamycin-induced DiCre-mediated recombination (**Fig. S1A, B**). Transgenic parasites were generated using a Cas9-mediated strategy to induce targeted double-stranded DNA breaks and eliminate non-recombinant parasites^19^. Following the selection of transgenic parasites, limiting dilution cloning was performed to isolate a genetically homogeneous line, designated PfDCX-HA:cKO. Correct modification of the *PfDCX* locus was confirmed by diagnostic PCR and Sanger sequencing (**Fig. S1C**).

PfDCX-HA could not be detected by western blot analysis of schizont-stage parasites (**Fig. 1A**), and no DCX signal was observed in merozoites by either immunofluorescence assay (IFA - **Fig. S1F**) or Ultrastructure Expansion Microscopy^20^ (U-ExM - **Fig. 1B**). To ensure that the fixation conditions did not compromise the integrity of the conoid complex, we performed a parallel analysis using parasites expressing C-terminally tagged MyoB-GFP, a known marker of the conoid complex in *P. berghei*^11^ that was shown to be expressed at the apex of *P. falciparum* schizont^21^. In *P. falciparum* MyoB-GFP schizonts, U-ExM revealed a distinct MyoB-GFP ring structure at the apex of merozoites, indicating that our experimental conditions preserve the architecture of the conoid complex and allow its visualisation in merozoites (**Fig. 1C**).

**Figure 1.**
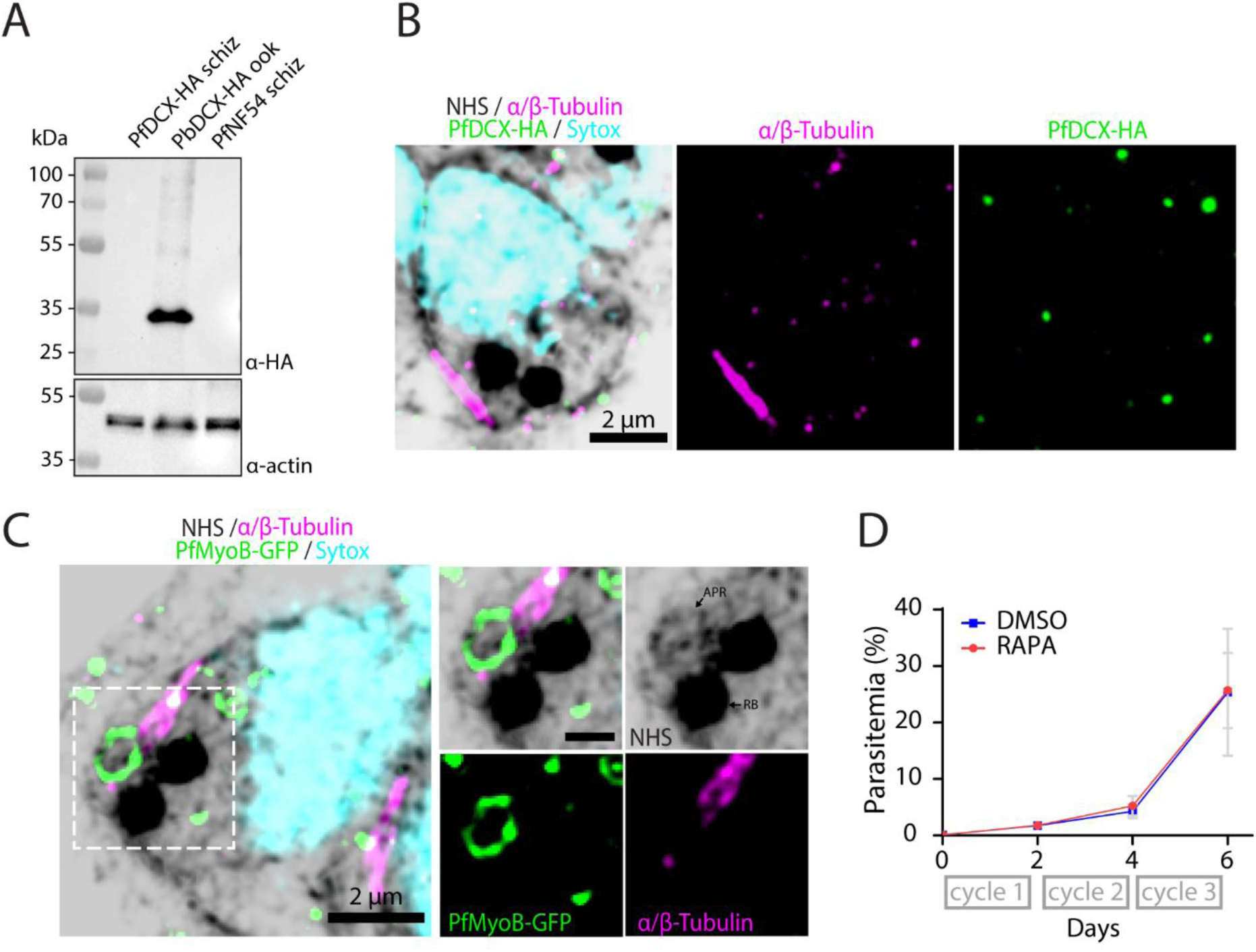
DCX is not expressed in *P. falciparum* merozoites and is dispensable for the proliferation of asexual blood stages. **(A)** DCX is not detectable in PfDCX-HA schizonts nor in the NF54Dicre parental line but is expressed in *P. berghei* PbDCX-HA ookinetes, as shown by immunoblotting probed with monoclonal α-HA antibodies (expected molecular weight: 33 kDa). α-Actin is shown as a loading control. **(B)** Representative U-ExM images of a PfDCX-HA merozoite showing DNA (sytox, cyan), rhoptries (NHS, black), subpellicular microtubules (α/β-Tubulin, magenta), and absence of specific detectable PfDCX-HA signal (green). Scale bar: 2 µm. **(C)** Representative U-ExM images of a *P. falciparum* merozoite expressing PfMyoB-GFP, which forms a ring at the apical pole. Shown are DNA (sytox, cyan), rhoptries (NHS, black) and SPMTs (α/β-Tubulin, magenta). APR, apical polar ring; RB, rhoptry bulb. Scale bars: 2 µm (left panel), 1 µm (insets). **(D)** Growth curve over three asexual replicative cycles of PfDCX-HA:cKO parasites treated with DMSO or rapamycin (RAPA), showing no detectable growth defect upon *PfDCX-HA* deletion. Data is representative of three biological replicates (mean ± SD, Šídák’s multiple-comparisons test).

To further investigate the role of *DCX* in asexual blood-stage proliferation, we assessed the phenotypic consequences of its conditional deletion. *PfDCX* was conditionally deleted by transient treatment with rapamycin using DMSO as a vehicle control. Successful excision of the *PfDCX* gene was confirmed by diagnostic PCR (**Fig. S1D-E**). Monitoring of PfDCX-HA:cKO parasites treated with either DMSO or rapamycin under shaking conditions over three replication cycles revealed that *PfDCX* deletion did not affect the proliferation of *P. falciparum* asexual blood stages (**Fig. 1D**).

### DCX is required for efficient transmission of *P. falciparum* to the mosquito vector

We next sought to evaluate the role of PfDCX in *P. falciparum* transmission to *Anopheles coluzzii* mosquitoes. To do so, PfDCX-HA:cKO gametocytes were induced *in vitro*, and either rapamycin or DMSO was added 24 hours post-induction to sexually committed ring-stage parasites for a duration of two days (**Fig. 2A**). Western blot analysis of stage III gametocytes revealed detectable PfDCX expression in DMSO-treated samples, while no signal was observed in rapamycin-treated parasites, further confirming successful gene deletion (**Fig. 2B**). Conditional deletion of *PfDCX* did not impair gametocyte maturation up to stage V, as assessed by microscopic examination of the perimeter to circularity ratio in Haemacolor-stained preparations (**Fig. 2C, D**).

**Figure 2.**
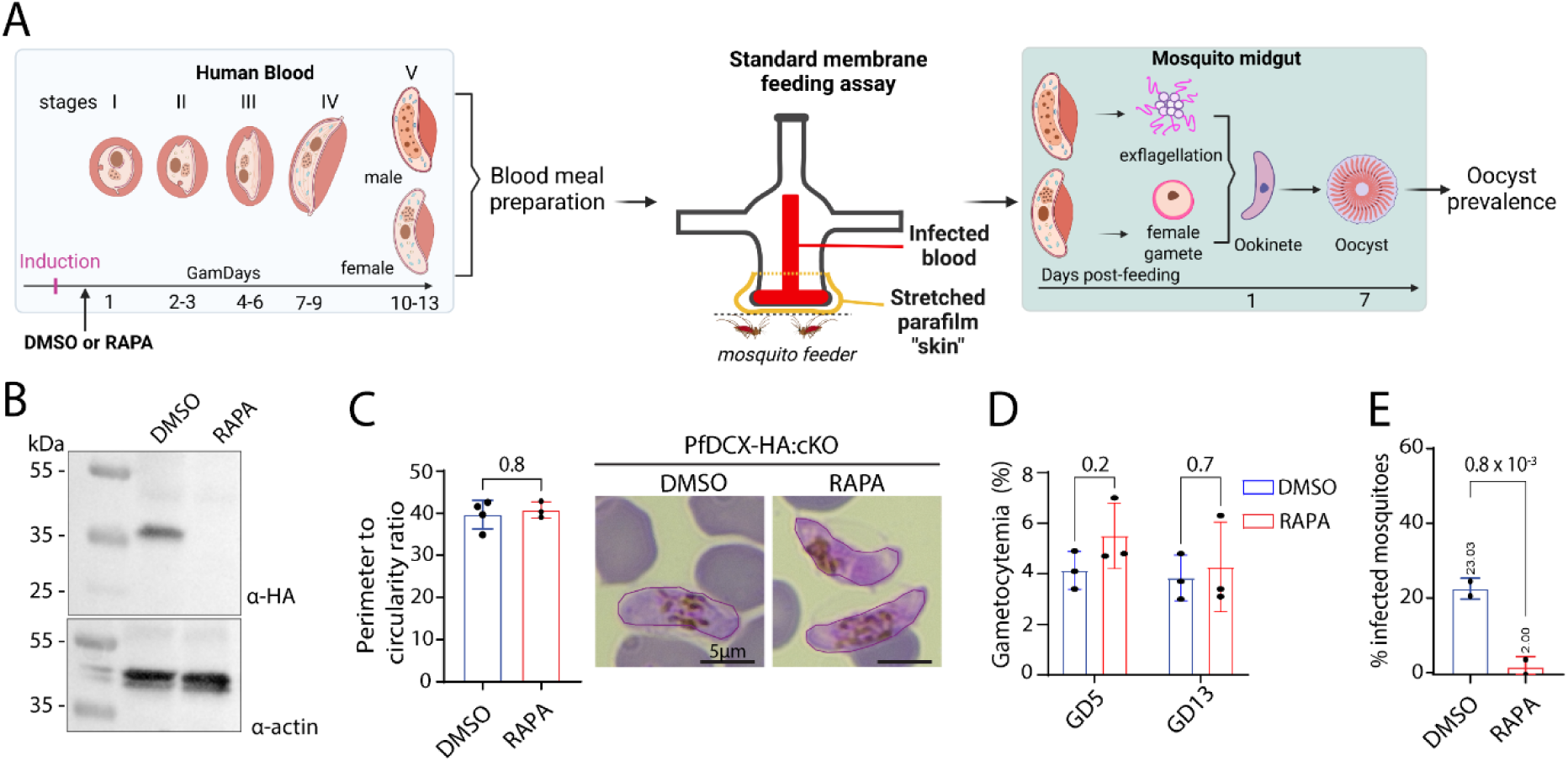
PfDCX is required for efficient mosquito infection by *P. falciparum*. **(A)** Schematic of the experimental workflow used to induce *P. falciparum* gametocytogenesis *in vitro* and to perform standard membrane feeding assays (SMFAs). Gametocytes were treated with DMSO or rapamycin one day after gametocyte induction. Gametocyte development proceeds from GamDay1 (stage I gametocytes) to full maturity at GamDay13 (stage V gametocytes). Blood meals prepared from gametocyte cultures were used to feed *Anopheles culozzii* mosquitoes at 37°C. Upon entry into the mosquito midgut, gametocytes are activated, and microgametes undergo exflagellation, while female gametocytes round up. Fertilisation leads to zygote formation, and mature ookinetes develop within 24 hours to further form oocysts. Seven days after the bloodmeal, mosquitoes were dissected, and the prevalence of oocysts was quantified. **(B)** Immunoblot showing the expression of PfDCX-HA in stage III gametocytes treated with DMSO, and efficient conditional depletion upon rapamycin treatment. Blots were probed with monoclonal α-HA antibodies; α-Actin serves as a loading control. **(C)** Left: quantification of stage V PfDCX-HA:cKO gametocyte shape, measured as the perimeter-to-circularity ratio on Haemacolour-stained cells, showing no significant difference between DMSO- and rapamycin-treated parasites (mean ± SD, DMSO n=4, RAPA n=3 biological replicates, paired two-tailed *t* test). Right: representative images of stage V PfDCX-HA:cKO gametocytes treated with DMSO or rapamycin, with cell perimeters outlined. Scale bar 5µm. **(D)** Deletion of *PfDCX* in gametocyte cultures does not affect gametocytaemia at day GamDay 5 or GamDay 13 (mean ± SD, n=3 biological replicates, 2-way ANOVA multiple-comparisons correction). **(E)** Oocyst prevalence is significantly reduced in mosquitoes infected with rapamycin-treated parasites compared with DMSO-treated controls (mean ± SD, n=2 biological replicates, paired two-tailed *t* test, 25 to 38 mosquitoes per condition per replicate).

We then investigated whether PfDCX is required for successful colonisation of the mosquito vector by assessing oocyst formation in the midgut following standard membrane feeding assays (**Fig. 2A**). Seven days post-infection, mosquito midguts were dissected, and oocysts were counted using brightfield microscopy. Oocyst prevalence was defined as the proportion of mosquitoes harbouring at least one oocyst. Parasites treated with rapamycin exhibited a significant 11-fold reduction in infection prevalence compared to the DMSO control, indicating that PfDCX is required for efficient transmission of *P. falciparum* to *Anopheles coluzzii* mosquitoes (**Fig. 2E and Fig. S2**).

### DCX forms a ring at the apex of *P. falciparum* ookinetes and co-localises with conoidal tubulin fibres in *P. berghei* ookinetes

To investigate the role of PfDCX during mosquito-stage development, we first attempted to localise PfDCX-HA in *in vitro* developing ookinetes as previously described^22^. Although we were unable to obtain fully formed, banana-shaped ookinetes, we observed retort forms displaying apical PfDCX-HA localisation in DMSO-treated parasites, suggesting a conoid complex localisation (**Fig. S3A**). Rapamycin-treated parasites also produced retort forms, but with a significantly thinner and elongated protrusion extending from the residual body compared to those observed in the DMSO control (**Fig. S3B, C**).

To examine PfDCX-HA localisation in fully formed ookinetes, we purified PfDCX-HA:cKO parasites from mosquito midguts of DMSO-treated controls and performed U-ExM analysis. The PfDCX-HA signals appeared as a ring structure at the apex of the cell (**Fig. 3A**). However, the limited number of cells recovered from mosquito midguts prevented us from definitively localising PfDCX to the conoid, as colocalisation with the conoid α- and β-Tubulin signals could not be reliably established.

**Figure 3.**
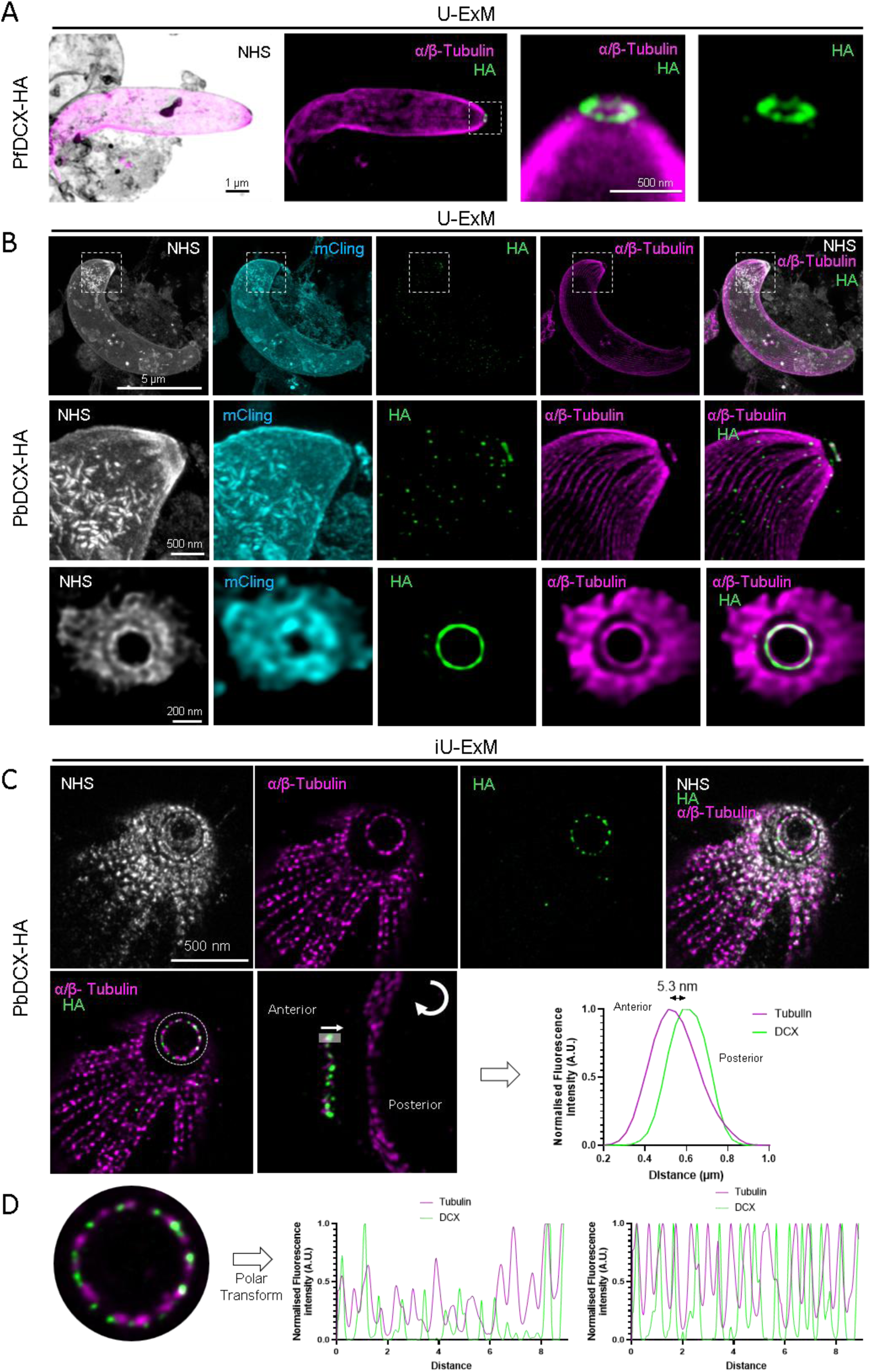
DCX associates with conoid tubulin fibres in both *P. falciparum* and *P. berghei* ookinetes. **(A)** Representative U-ExM images of a *P. falciparum* ookinete expressing PfDCX-HA purified from mosquito midguts stained for α/β-Tubulin (magenta) and PfDCX-HA (green). **(B)** U-ExM representative images of an *in vitro*-produced *P. berghei* ookinete expressing PbDCX-HA colocalising with the tubulin-based conoid. The first two lanes show a longitudinal view of an ookinete with insets highlighting the apical complex stained with NHS-ester (protein density in white), mCling (membranes in cyan), HA (PbDCX-HA in green) and α/β-Tubulin (magenta). The third lane shows a top view of the apical complex. **(C)** Top panels: Representative iterative U-ExM images of a *P. berghei* ookinete expressing PbDCX-HA, allowing resolution of separated conoid tubulin fibres stained with α/β-Tubulin (magenta) and interleaved with PbDCX-HA (green). The NHS-ester staining highlights protein density. Bottom panels: top and side views of the ookinete conoid showing PbDCX-HA localisation along tubulin fibres, excluding their apical ends (left and middle panels). Normalised fluorescence intensity of PbDCX and α/β-Tubulin. The fluorescence intensity peaks are separated by 5.3nm (right panel). **(D)** Top view of PbDCX-HA and tubulin fibres. Polar transformation of the corresponding fluorescence signals is shown without (left) and with normalisation (right), illustrating the alternation between the two signals.

To address this limitation, we employed the *P. berghei* model, which enables the *in vitro* production of a large number of motile ookinetes. We generated a transgenic *P. berghei* line in which *DCX* was endogenously tagged with a C-terminal 3xHA tag. Successful integration was confirmed by diagnostic PCR (**Fig. S4A-B**) and western blot analysis (**Fig. 1A**). Using U-ExM, we observed early expression of PbDCX-HA before the transition from the round zygote to the developing ookinete (**Fig. S3D**). In banana-shaped ookinetes, PbDCX-HA localised to the conoid complex, as revealed by co-staining with α- and β-tubulin to mark microtubules, NHS-ester to visualise protein density, and mCling to delineate membranes (**Fig. 3B**).

To further refine the precise localisation of PbDCX-HA within the conoid complex, we applied iterative Ultrastructure Expansion Microscopy (iU-ExM), which enables a 14- to 20-fold physical expansion of biological samples^23^. The increased resolution afforded by iU-ExM allowed individual tubulin fibres within the conoid to be resolved, together with concentric ring-like structures revealed by NHS-ester staining that likely correspond to the APR. To determine the precise position of PbDCX-HA within the conoid, the individual structure was aligned along its anterior–posterior axis and analysed by fluorescence intensity profiling. This revealed that the PbDCX-HA signal peaked approximately 5 nm posterior to the distal ends of the conoid fibers. In addition, analysis of *en face* views showed that PbDCX-HA was laterally shifted relative to the tubulin fibres rather than fully colocalising with them. Together, these data indicate that DCX occupies a distinct subdomain of the conoid rather than the microtubule tips or walls (**Fig. 3C and 3D**). This localisation is consistent with DCX decorating the inter-fibre crests of conoid microtubules, as observed in *T. gondii* [14], thereby supporting a role for DCX in organising the ookinete conoid architecture in *Plasmodium*.

### PfDCX is required for microgamete formation

Given our observation of DCX expression in *P. falciparum* gametocytes, we hypothesised that its requirement for mosquito transmission might be linked to an earlier role during sexual development. During microgametogenesis, the component parts of eight axonemes are assembled within the cell cytoplasm to form eight flagellated male gametes in a process called exflagellation, which is easily quantifiable by light microscopy. Gametogenesis was induced in stage V PfDCX-HA:cKO gametocytes treated with DMSO or rapamycin, and exflagellation centres were counted 15 min following reduction in temperature and gametocyte activation with the mosquito-derived small molecule xanthurenic acid. Conditional deletion of *PfDCX* led to a significant 3-fold reduction in exflagellation rate, indicating a role of DCX prior to fertilisation and ookinete formation (**Fig. 4A**).

**Figure 4.**
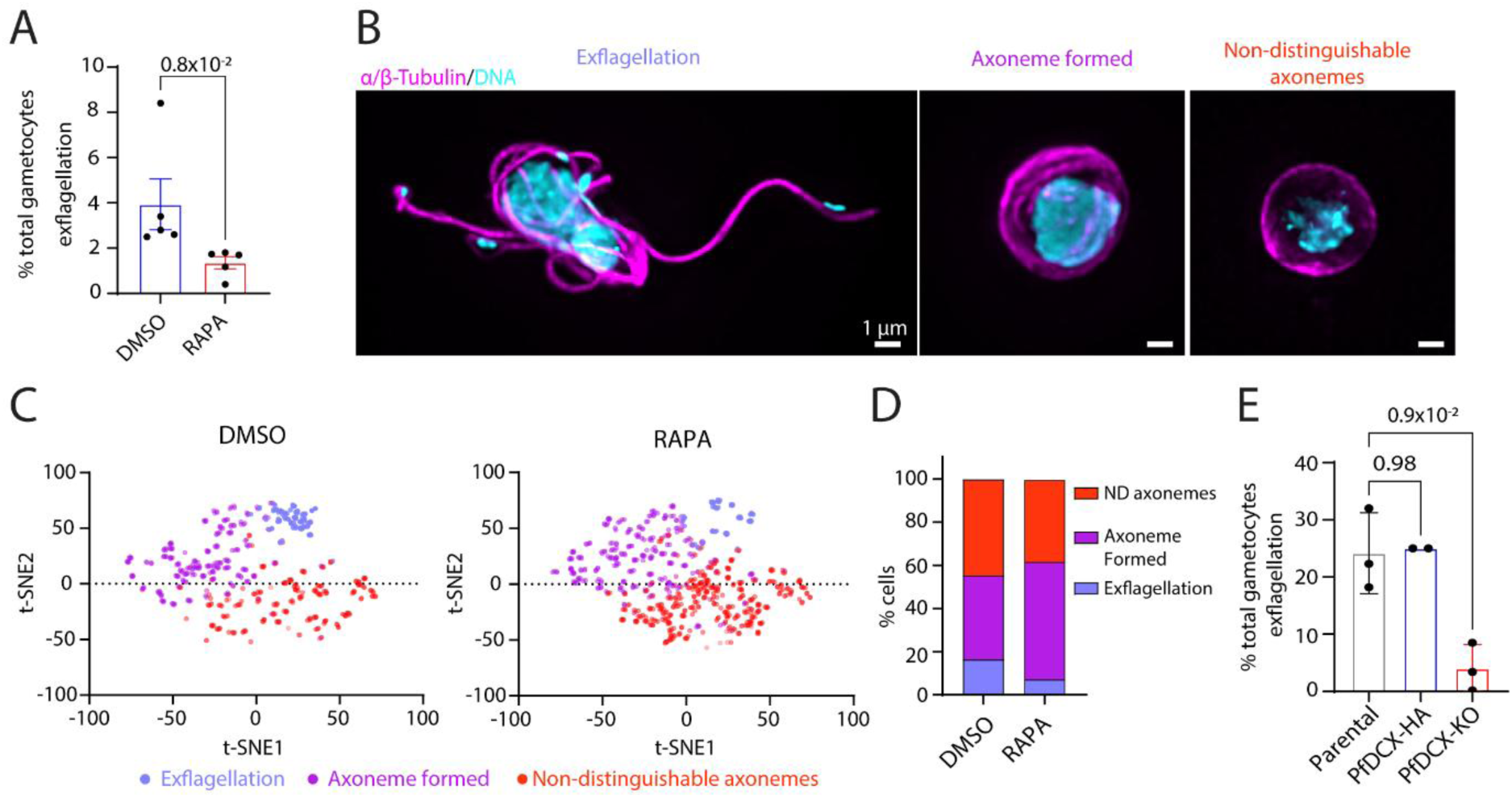
Microgametogenesis is affected in PfDCX-cKO parasites. **(A)** Exflagellation is reduced by 3-fold in rapamycin-treated PfDCX-HA:cKO parasites following activation with XA compared to DMSO-treated controls as assessed by live microscopy (mean ± SD, n=5 biological replicates, Mann-Whitney test). **(B)** A total of 2’680 cells from DMSO- and rapamycin-treated conditions were randomly imaged, processed, and clustered using the PhIDDLI pipeline using K-means clustering. Representative immunofluorescence images for each cluster are shown with exflagellating microgametes (left), cells containing assembled cytosolic axonemes (centre), and cells with non-distinguishable (ND) axonemes (right). Axonemes are labelled with α/β-Tubulin (magenta) and DNA with Hoechst (cyan). Scale bar: 1 µm. **(C-D)** *PfDCX* deletion in PfDCX-HA:cKO gametocytes treated with rapamycin results in a reduced proportion of exflagellating microgametes and mirrored by an increase in cells with formed axonemes relative to DMSO-treated parasites. **(E)** Exflagellation is reduced by 6-fold in a clonal PfDCX-KO parasite line compared to the NF54Dicre line and the parental PfDCX-HA:cKO lines grown in the absence of DMSO (mean ± SD, n=3 biological replicates, one-way ANOVA).

To determine whether this reduction was due to a decrease in male gametocyte numbers, we assessed the sex ratio of gametocytes upon *PfDCX* deletion. This was monitored by Pfg377 expression in stage V gametocytes, as Pfg377 is associated with osmiophilic bodies predominantly found in *P. falciparum* female gametocytes^24^. Conditional deletion of *PfDCX* led to a minor 1.4-fold decrease in microgametocytes (**Fig. S5A, B**), which does not account for the observed reduction in exflagellation. To characterise the stage of exflagellation affected in PfDCX-cKO male gametes, a pre-trained machine learning image analysis pipeline (PhIDDLI) was applied to immunofluorescent images of activated microgametes as described previously^25^. Using k-means clustering, cells were grouped into three clusters, which were separated based on their morphological phenotypes. The first cluster comprised cells that had successfully undergone exflagellation, characterised by clearly formed flagella. The second cluster included cells displaying cytoplasmic axoneme assembly without exflagellation, while the third cluster consisted of microgametocytes showing no detectable axoneme formation (**Fig. 4B**). Parasites lacking *PfDCX* exhibited a reduced proportion of exflagellating cells, confirming the observed exflagellation defect. This decrease was mirrored by a corresponding increase in gametocytes that assembled axonemes but failed to undergo exflagellation. In contrast, the proportion of parasites lacking detectable axonemes was similar under both conditions, indicating that deletion of *PfDCX* does not impair axoneme assembly *per se*, as assessed by IFA (**Fig. 4C, D**).

To rule out the possibility that residual exflagellation centres observed in rapamycin-treated cells were linked to incomplete excision of the *PfDCX* locus, we generated a PfDCX-KO line by cloning PfDCX-HA:cKO asexual blood stages following a rapamycin treatment for 48 h. Exflagellation was further reduced by 6.2-fold in knockout parasites relative to the non-excised PfDCX-HA–tagged parental line, indicating that PfDCX contributes to efficient exflagellation but is not strictly essential (**Fig. 4E**).

### PfDCX is required for axonemal symmetry during microgametogenesis

We next sought to refine our understanding of the cellular defects underlying impaired exflagellation by first focusing on the microtubule-based axonemes using U-ExM on non-activated gametocytes and gametes activated for 15 minutes, labelled with α- and β-tubulin (**Fig. 5A**). Consistent with live microscopy observations, PfDCX-KO parasites exhibited a reduced number of exflagellating microgametes compared to control cells (**Fig. 5B**).

**Figure 5.**
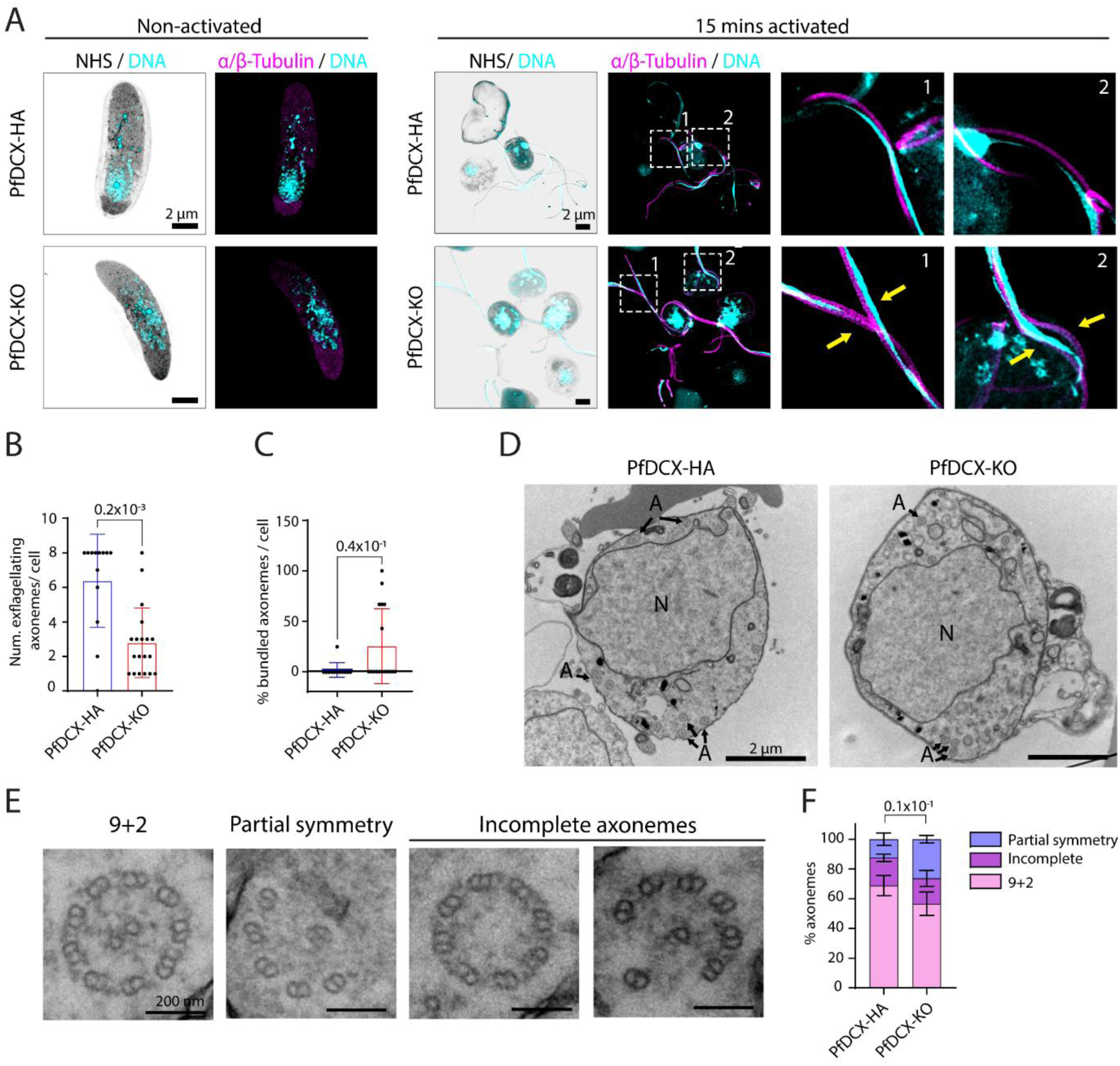
PfDCX-KO microgametes show altered axonemal symmetry. **(A)** Representative U-ExM images of PfDCX-HA and PfDCX-KO non-activated stage V gametocytes (left), and microgametes 15 minutes post-activation (right). Yellow arrowheads indicate bundled axonemes observed in DCX-KO activated microgametes. Scale bars: 2 µm. **(B)** Quantification from U-ExM images of the number of exflagellating axonemes emerging from the residual body, confirming a significant reduction in PfDCX-KO microgametes (mean ± SD, n = 2 biological replicates, unpaired *t*-test). **(C)** Increased proportion of bundled axonemes per cell in PfDCX-KO microgametes (mean ± SD, n = 2 biological replicates, unpaired *t*-test). **(D)** Representative transmission electron microscopy (TEM) images of PfDCX-HA and PfDCX-KO exflagellating microgametocytes, 15 minutes post-activation. Black arrows indicate axonemes. A, axoneme; N, nucleus. Scale bar: 2 µm. **(E)** Representative TEM images illustrating the three axonemal microtubule organisations used for classification: canonical 9+2 conformation, axonemes with partial symmetry, and incomplete axonemes, which lack the central pair or one or more doublets. Scale bar: 200 nm. **(F)** PfDCX-KO parasites display a reduced proportion of axonemes with the 9+2 architecture and an increased proportion of axonemes with partial symmetry. Quantification was performed on TEM sections, analysing 93 axonemes from 37 cells for PfDCX-HA and 236 axonemes from 54 cells for PfDCX-KO. (mean ± SD, n = 2 biological replicates; χ² test with 2 degrees of freedom).

Interestingly, U-ExM enabled quantification of a higher proportion of bundled axonemes in the remaining exflagellating microgametes of PfDCX-KO parasites (**Fig. 5C**), suggesting defective axoneme assembly. To further investigate axonemal architecture, we performed transmission electron microscopy (TEM), categorising axonemes into three structural types: (i) canonical 9+2 organisation with nine microtubule doublets and a central pair of singlets; (ii) partial symmetry; (iii) incomplete axonemes, lacking the central pair or missing one or more doublets (**Fig. 5D-E**). PfDCX-KO parasites showed a decreased proportion of axonemes with a canonical 9+2 structure, mirrored by a modest but significant increase in partially symmetric axonemes (**Fig. 5F**), indicating that PfDCX is modestly required for proper axoneme assembly and symmetry during microgametogenesis.

### PfDCX is required for branching of subpellicular microtubules in stage IV gametocytes

Although axoneme bundling and a reduction in axonemal symmetry were observed in developing microgametes, these defects are relatively modest and are unlikely to solely explain the observed decrease in exflagellation rate. As we noted expression of PfDCX-HA early in gametocytogenesis, we next wondered whether the protein could be important for the atypical organisation of gametocyte SPMTs. U-ExM did not allow to detect differences in overall SPMT organisation in stage IV gametocytes, which retained their characteristic falciform shape as highlighted by NHS-ester staining and displayed SPMTs stained with α/β-Tubulin (**Fig. 6A**). To further gain in resolution, we analysed stage IV gametocytes by TEM, which uncovered altered SPMT organisation in PfDCX-KO parasites. While control PfDCX-HA parasites predominantly displayed SPMTs branched into doublet, triplet, and occasional quadruplet structures, PfDCX-KO parasites exhibited a significantly higher proportion of single microtubules (**Fig. 6B and C**). However, this reduction in microtubule branching was not associated with a reduced linear density of SPMTs below the IMC (**Fig. 6D**), suggesting that PfDCX requirement is for stabilising branched SPMTs. While DCX-HA expression was detected by western blot in gametocytes, we were unable to localise the protein using either IFA or U-ExM possibly due to low or transient expression, which may prevent its detection with our current approaches.

**Figure 6.**
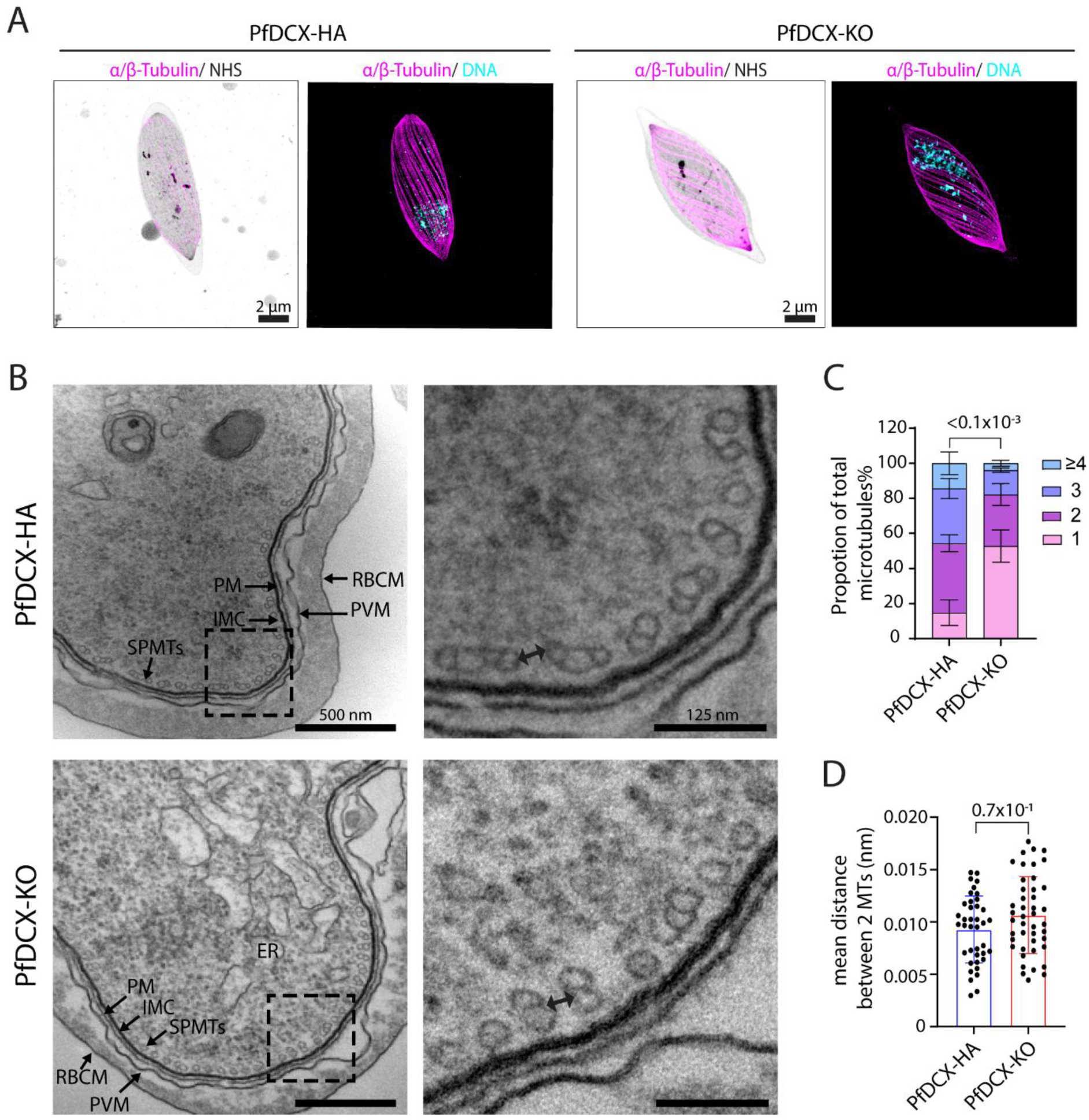
PfDCX participates to branching of subpellicular microtubules in stage IV gametocytes. **(A)** Representative U-ExM images of PfDCX-HA and PfDCX-KO stage IV gametocytes displaying their characteristic elongated morphology supported by SPMTs (α/β-Tubulin, magenta; NHS ester, grey scale). Scale bar: 2 µm. **(B)** Representative TEM images of cross-sections through PfDCX-HA and PfDCX-KO stage IV gametocytes. Black double-headed arrows indicate the measured distance between groups of branched SPMTs. RBCM, red-blood cell membrane; PVM, parasitophorous vacuole membrane; PM, plasma membrane; IMC, inner membrane complex; ER, endoplasmic reticulum. Scale bars: 500 nm, inset: 125 nm. **(C)** PfDCX-KO stage IV gametocytes exhibit an increased proportion of single microtubules (1) compared with PfDCX-HA gametocytes, which have a majority of double and triple MTs (2, 3). Quantification was performed on TEM sections, analysing 49 cells for PfDCX-HA and 85 cells for PfDCX-KO (mean ± SD, n=2 biological replicates; χ² test with 2 degrees of freedom). **(D)** Quantification of the SPMT linear density beneath the IMC, expressed as the mean distance between two SPMTs in nanometres, shows no difference between PfDCX-HA and PfDCX-KO stage IV gametocytes (mean ± SD, n=2 biological replicates, unpaired *t*-test).

### SPMTs in stage IV micro- and macrogametocytes display distinct levels of polyglutamylation independent of DCX

It was recently shown that SPMTs become polyglutamylated in stage IV gametocytes, potentially contributing to the stabilisation and regulation of the microtubule network^26^. In human cells, DCX has been shown to restrict neuronal branching by regulating tubulin polyglutamylation^27^. We therefore asked whether an interplay between DCX and tubulin polyglutamylation might similarly regulate SPMT branching in *P. falciparum* gametocytes.

We first examined tubulin polyglutamylation in the parental line. Unexpectedly, we observed previously unreported differences in polyglutamylation levels between macro- and microgametocytes (**Fig. 7A**). Macrogametocytes, identified by the high density of osmiophilic bodies following NHS-ester staining, showed a strong SPMT polyglutamylation signal as assessed by corrected total cell fluorescence. In contrast, microgametocytes characterised by their NHS-ester-dense bipartite MTOC^28^ displayed markedly lower polyglutamylation signals, despite comparable detection of α- and β-Tubulin. However, deletion of *DCX* did not lead to significant changes in polyglutamylation patterns, suggesting that the DCX-dependent differences in SPMT branching are not linked to tubulin polyglutamylation (**Fig. 7B and C**).

**Figure 7.**
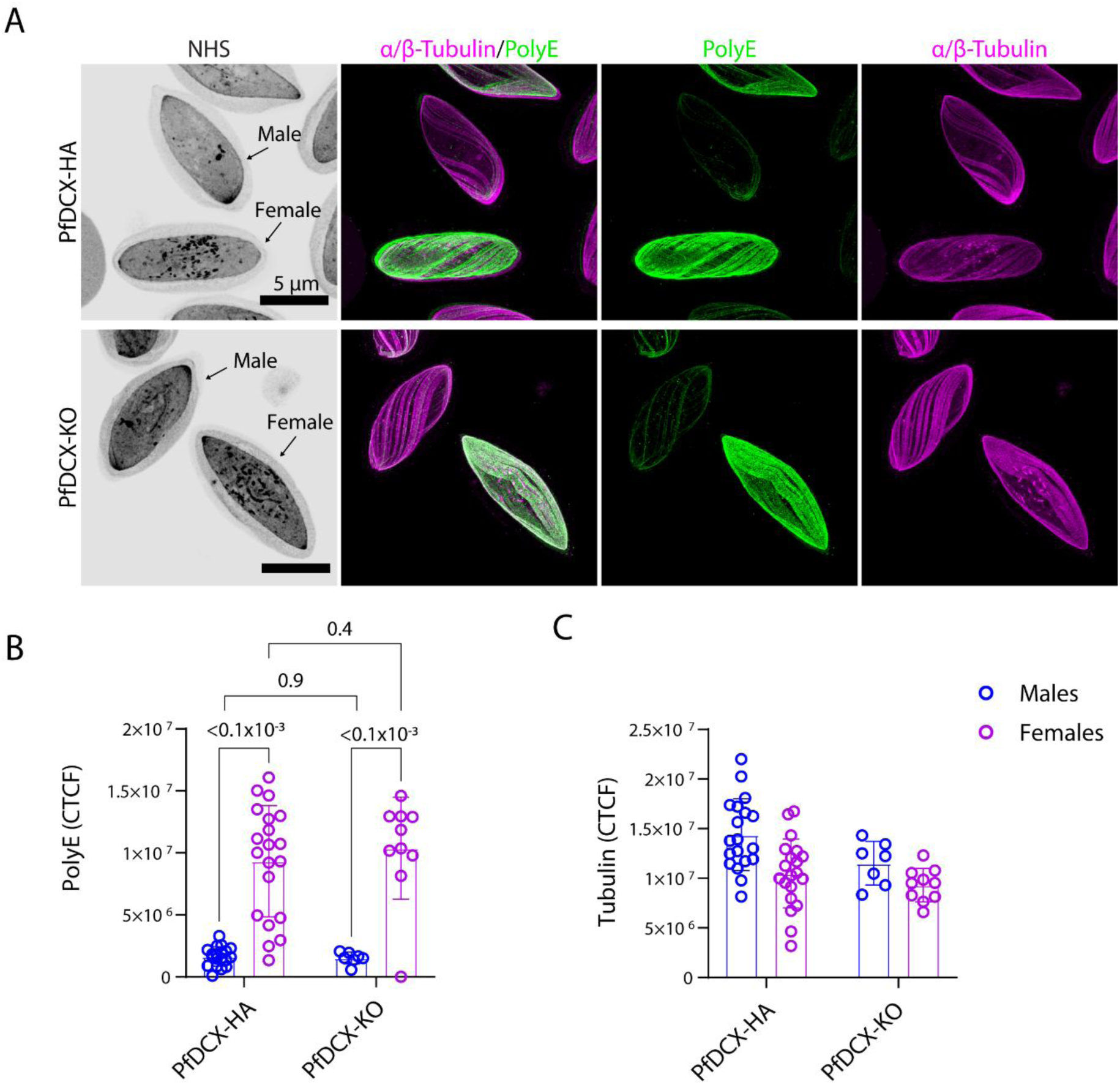
SPMTs in stage IV micro- and macrogametocytes display distinct levels of polyglutamylation independent of DCX. **(A)** Representative U-ExM images of PfDCX-HA and PfDCX-KO stage IV gametocytes displaying their characteristic elongated morphology supported by SPMTs (α/β-Tubulin, magenta). NHS-ester staining (black) highlights the high density of osmiophilic bodies in macrogametocytes (females) and the large bipartite MTOC of microgametocytes (males). Staining for polyglutamylation (PolyE, green) reveals differential signals between macro and microgametocytes. Scale bar: 5 µm. **(B-C)** Quantification of the PolyE (B) and α/β-Tubulin (C) signals assessed by corrected total cell fluorescence (CTCF) in macro- and microgametocytes of the DCX-HA and DCX-KO lines.

### *PfDCX* deletion increases protofilament numbers in SPMTs

Given that DCX has previously been implicated in the stabilisation of microtubule protofilaments in other biological systems^13,15^, we investigated its role by examining the molecular organisation of protofilaments of SPMTs in stage IV gametocytes. To assess the impact of PfDCX on protofilament architecture within the SPMTs, we performed in situ cryo-electron tomography (CryoET) on stage IV gametocytes (**Fig. 8A & B**).

**Figure 8.**
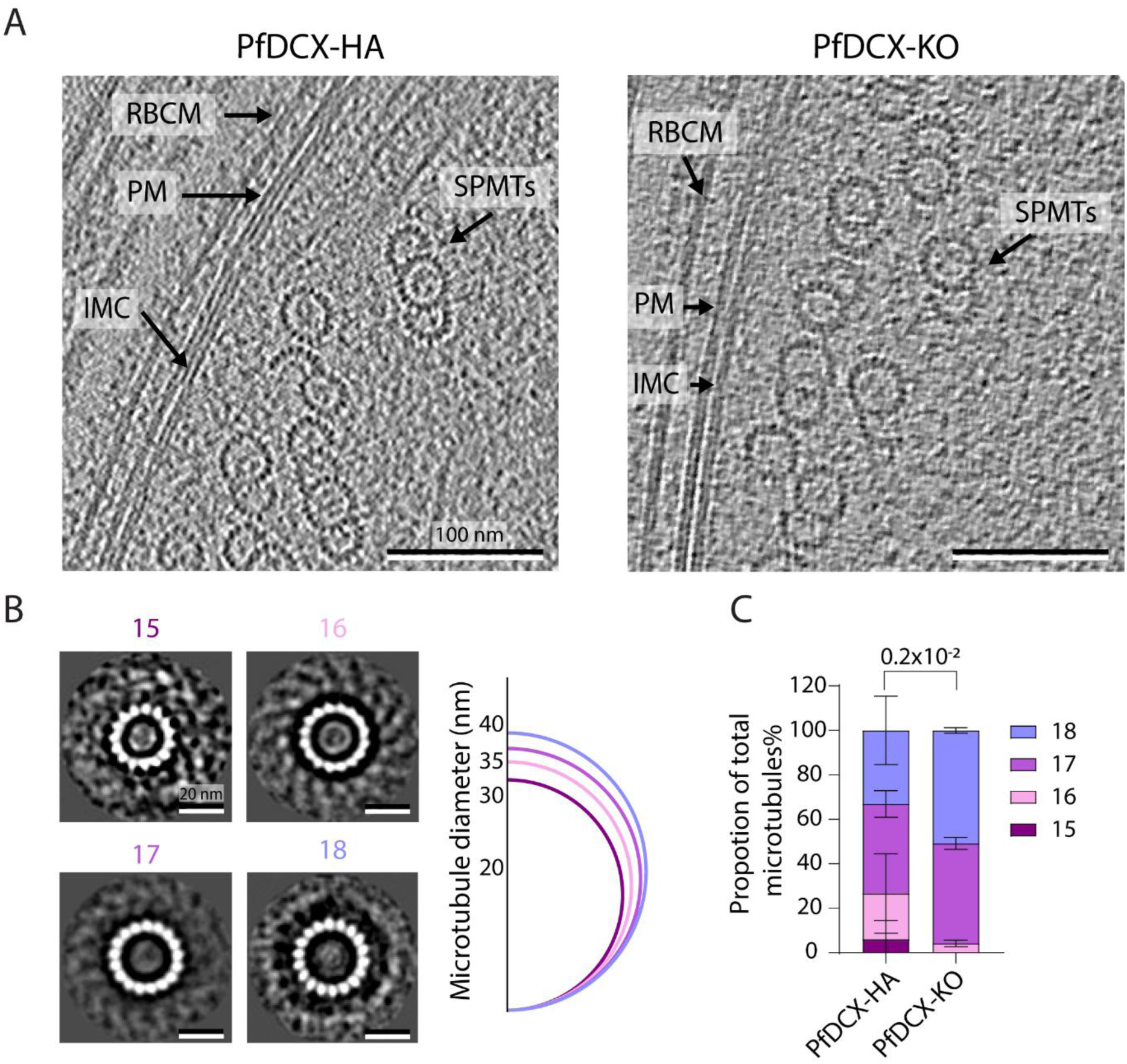
PfDCX contributes to the 13- to 18-protofilament architecture of subpellicular microtubules in stage IV gametocytes. **(A)** Representative cryo-electron tomography (Cryo-ET) images through PfDCX-HA and PfDCX-KO stage IV gametocytes. RBCM, red-blood cell membrane; PM, plasma membrane; IMC, inner membrane complex. Scale bar: 100 nm**. (B)** Representative 2D projections of subtomogram averages of single microtubule containing 15, 16, 17 or 18 protofilaments. The right panel shows the average microtubule diameter for each protofilament category. No 13-and 14-protofilament microtubules were observed in these samples. Scale bar: 20 nm. **(C)** Distribution of protofilament numbers in SPMTs of stage IV gametocytes from the PfDCX-HA (n=43 microtubules) and PfDCX-KO (n=90 microtubules) parasite lines. The distribution differs significantly with PfDCX-KO parasites enriched for higher protofilament numbers (17-18) relative to PfDCX-HA (mean ± SD, n=2 biological replicates, Fisher’s exact test, Freeman–Halton extension, effect size Cramér’s V = 0.33).

The SPMTs that were imaged show a notable diversity in protofilament number and the presence of partial doublet, triplet, and quadruplet structures, in agreement with the literature^4^. These non-canonical microtubule geometries were seen in both PfDCX-HA and PfDCX-KO cells, and no microtubules were seen with fewer than 15 protofilaments in either condition. To quantify these geometries, per-microtubule subtomogram averages were acquired, and the protofilament number was determined by applying symmetry-based methods using custom scripts. Statistical analysis, specifically a Fischer’s exact test, determined that DCX knockout results in an enrichment of microtubules of greater numbers of protofilaments, with a markedly reduced proportion of microtubules containing fewer than 17 protofilaments, and no microtubules containing fewer than 16 (**Fig. 8C**). Thus, it can be concluded that DCX plays a role in the observed diversity in microtubule architecture in gametocytes.

## Discussion

In this study, we identified an unexpected cellular requirement for DCX that is critical for the transmission of *P. falciparum* to the mosquito vector. In apicomplexan parasites, DCX function has so far been associated exclusively with the conoid complex, a specialised parasitic structure tailored for motility and host cell invasion^6^. In *T. gondii*, the conoid is composed of up to 14 tightly curved and tilted tubulin fibres, each consisting of nine protofilaments, forming a comma-shaped strip in cross-section rather than a hollow tube. These structural properties have been attributed to tubulin-binding proteins, including DCX^16,29^, which localises specifically to the conoid tubulin fibers and contributes to its stability^13^. Remarkably, *Toxoplasma* DCX has been shown to stabilise curved and open microtubules in heterologous systems^15^. Despite its conserved association with conoid fibres in *P. berghei* ookinetes, deletion of *DCX* in this species did not impair either ookinete gliding motility or transmission through the mosquito^14^, leaving its molecular role in the *Plasmodium* conoid unresolved. Microtubule fibres within the ookinete conoid have been shown to adopt a comma-shaped configuration^4^, suggesting a DCX-dependent scaffolding function similar to that observed in *T. gondii*. Here, we demonstrate that DCX localises along conoid fibres in ookinetes, likely on their crest, as observed in *T. gondii*^23^. In the absence of a clear ookinete phenotype associated with *DCX* deletion in *P. berghei*^14^, it is plausible that other conoidal proteins compensate for the loss of DCX in *Plasmodium*. In *Toxoplasma*, DCX has been shown to form a meshwork over protofilaments 5–9 of microtubule fibres together with CPH1 and CF6^16^. While CF6 does not appear to be conserved in *Plasmodium*, CPH1 is conserved and its expression peaks in ookinetes, raising the possibility that CPH1 may compensate for *DCX* deletion. It is also possible that the reduced structural complexity of the ookinete conoid renders it less reliant on microtubule stability for its function.

While we cannot rule out a species-specific requirement for DCX in *P. falciparum* ookinete motility linked to conoid function, our findings reveal a notable species-specific difference with DCX being essential for efficient transmission of *P. falciparum* to the mosquito vector. We attribute this transmission requirement to a role for DCX in non-motile intracellular sexual stages beyond its function at the conoid. Specifically, we found that DCX is essential for maintaining the unique architecture of gametocyte SPMTs by promoting the branching of open microtubules. *DCX* deletion led to a reduction in the number of microtubules per raft, while increasing the number of protofilaments per remaining microtubule. Interestingly, the density of microtubule rafts beneath the inner membrane complex remained unaffected by DCX loss. We therefore propose that DCX facilitates the opening or stabilisation of open microtubules, enabling branching around a closed microtubule. Interestingly, although DCX localises to the conoid and is required for the proper organisation of gametocyte SPMTs, it does not appear to be necessary for SPMT function in either ookinetes or merozoites, which exhibit more conventional 13-protofilament microtubules^4^. This differential localisation suggests that other MAPs may play a key role in guiding DCX to specific microtubule populations and in contributing to their stage-specific requirement for DCX. Notably, although *PfDCX* deletion affects gametocyte SPMTs, the resulting phenotype on microtubule branching is significant yet not absolute, as branching is still observed upon *DCX* deletion. This further supports the idea that additional MAPs or factors are likely involved in the atypical organisation of SPMTs in *P. falciparum* gametocytes. For example, *Plasmodium*-specific sequences in tubulin isoforms alter the tubulin dimer structure, producing smaller but stronger lateral contacts and a stiffer lattice than those observed in brain microtubules or in microtubules with higher protofilament numbers *in vitro*^30^.

We observed additional developmental defects associated with early *DCX* deletion in gametocytes, including reduced microgamete formation associated with impaired axoneme assembly and altered symmetry likely due to the loss of microtubule doublets. Although DCX may contribute specifically to the stabilisation of microtubule doublets in *P. falciparum*, no exflagellation phenotype was reported in *P. berghei*, suggesting that the defects observed in *P. falciparum* may be indirect and linked to earlier interaction of DCX with tubulin prior to axoneme assembly. This kind of species specificity linked to the different requirements for SPMTs in gametocytes has previously been reported for the SPMT-associated protein SPM3, which localises to SPMTs in *P. falciparum* gametocytes. While *SPM3* deletion led to aberrant sexual development in *P. falciparum*, it had no impact on sexual development in *P. berghei*^31^. Similarly, while both PbDCX and PfDCX are expressed early during ookinete development, no developmental defects were observed in *P. berghei* ookinetes, either *in vitro* or *in vivo*, in contrast to the impaired development seen in *in vitro* cultured *P. falciparum* zygotes. However, given that current *in vitro* conditions do not support complete ookinete maturation, we cannot conclusively determine whether DCX plays a relevant role in *P. falciparum* ookinete development.

Altogether, since *P. berghei* gametocytes lack SPMTs, the differing requirement for DCX in transmission between *P. falciparum* and *P. berghei* is likely attributable to its specific role in organising gametocyte SPMTs in *P. falciparum*. This observation raises questions about the cellular and molecular roles of SPMTs in *P. falciparum* gametocytes. SPMTs provide the characteristic elongated, falciform shape of gametocytes and possibly contribute to cellular rigidity and mechanical stability, thereby enabling gametocytes to withstand mechanical stress^32^. However, because gametocyte shape and rigidity are not strongly constrained *in vitro*, the atypical architecture of *P. falciparum* gametocyte SPMTs may serve functions beyond mechanical support. This hypothesis is further supported by the distinct polyglutamylation patterns of SPMTs observed in macro- and microgametocytes, suggesting divergent functional requirements for SPMTs between the two gametocyte sexes. These findings open exciting avenues for future research into the biology and functional specialisation of *Plasmodium* SPMTs.

## Material and Methods

### *P. falciparum* maintenance and synchronisation

Lines were maintained *in vitro* in 5% haematocrit, in complete RPMI (Gibco, lot 2436708) medium containing AlbuMAX II (0.5%; Gibco), gentamicin (40 µg/ml; Gibco), hypoxanthine (200 µM; Sigma) and choline chloride (2 mM; Sigma). Human RBCs (obtained from the Bern Transfusion Centre) were used to grow parasites, which were maintained at 37°C, under shaking conditions, in a specific gas mixture (3% O2, 4% CO2 in N2). Cultures were routinely microscopically examined using Haemacolour-stained thin blood films (Sigma-Aldrich). To obtain highly synchronised parasites, ring stages were synchronised with a 5% sorbitol treatment (Sigma-Aldrich, #SLBJ4425V) for 5 minutes at 37°C, then washed and resuspended in complete RPMI medium. Schizont-stage parasites were isolated by gradient centrifugation over 63% (v/v) isotonic Percoll (GE Healthcare, Life Sciences) cushions and cultured in fresh RBCs, allowing merozoite invasion.

### Generation of DNA-targeting constructs

The *PfDCX-HA*:cKO/DiCre transgenic line was generated by modifying the endogenous *PfDCX* locus in the NF54/DiCre-expressing *P. falciparum* clone using the Cas9-mediated genome editing strategy. The plasmid pUC57 vector used was commercially synthesised (Genscript); the full validated DNA sequence is provided in **Data S1A**. The following elements are included in the targeting pUC57 vector: (i) a 5′ homology region 1 (HR1) of 1008 base pairs (bp). (ii) A recodonised *PfDCX* sequence with a *loxP*int module followed by a 6HA tag and a *Lox66* site at the 3′ end of the recodonised gene. (iii) A 3’ HR2 of 1008 bp (**Data S1B**). The final targeting repair plasmid was linearised with NotI overnight before transfection. To target the *PfDCX* locus, two guide RNA–encoding sequences were independently inserted into the pDC2-Cas9-hDHFR (human dihydrofolate reductase)/yFCU (yeast cytosine deaminase/uridyl phosphoribosyl transferase–containing plasmid)^33^ using primer pairs PfDCX gRNA5 (F5/R5) and PfDCX gRNA6 (F6/R6) (**Table S1**). Constructs were transfected into the *P. falciparum* NF54/DiCre line^34^. The oligonucleotides used to generate and genotype the tagged parasite lines are in **Table S1**.

### Transfection and selection of *P. falciparum* parasites

Parasites were synchronised by purifying schizont stages using a 63% Percoll gradient. All lines generated were derived from NF54/Dicre parasites. Transfections were performed by electroporation. For the *PfDCX-HA*:cKO/DiCre line: parasites were transfected with 50µg of donor plasmid and 50µg of gRNA-pDC2-CRISPR/Cas9 plasmids in Cytomix (120mM KCI, 0. 15mM CaCl2, 2mM EGTA, 5mM MgCI2, 10mM K2HPO4/KH2PO4, pH 7.6, 25mM Hepes, pH 7.6). Electroporation was performed using the GenePulser Xcell BioRad machine (310V, 950 µF, Infinite Ω, 2mm cuvette). Drug pressure was applied the following day by adding 5 nM WR99210 for 6 days. Cultures were maintained in the absence of drug pressure until a stable parasite population was obtained. Successful gene editing was confirmed by PCR on genomic DNA (gDNA) (**Fig. S1**). All oligonucleotides used for diagnostic integration PCRs are listed in **Table S1**.

The *PfDCX-KO* transgenic line was generated by treating *PfDCX-HA*:cKO/DiCre parasites with 100nM rapamycin for 48 h and performing limiting dilution cloning to isolate knocked-out clonal parasites.

### Selection of *P. falciparum* clones by limiting dilution cloning

Synchronous parasites at ring stage were diluted to two desired densities (0.5 parasites/200 µl and 0.1 parasites/200 µl) in Albumax-complemented medium with human RBCs at a haematocrit of 1%, then dispensed into flat-bottomed 96-well plates (Corning 3595) using 200 µl of culture per well. Plaques were formed after 11 days of incubation at 37°C, collected and expanded in complete Albumax medium at 5% haematocrit.

### Generation of *P. berghei* DCX Cas9 vector

Guide RNAs (gRNAs) were designed using the EuPaGDT platform. Candidate guides were ranked based on three main parameters: (i) the overall score assigned by the software, (ii) their distance from the intended editing site, and (iii) predicted off-target activity within the *Plasmodium berghei* ANKA genome (PlasmoDB version 28). Gene-coding sequences, together with their flanking 5′ and 3′ untranslated regions, were obtained from PlasmoDB. For vector construction, the selected gRNAs were inserted into pYCm_Pbu6-hdhfr/yfcu-Cas9 backbones digested with BsmBI. To enable directional cloning, pairs of complementary oligonucleotides (synthesised by Microsynth) were generated, each containing the gRNA sequence plus four-base overhangs compatible with the BsmBI restriction sites. Equal amounts of the oligonucleotides were phosphorylated with T4 polynucleotide kinase (NEB), annealed by incubation at 95 °C for 5 minutes, and gradually cooled at 5 °C per minute until 25 °C. The resulting double-stranded products were diluted (1:200) and ligated into the BsmBI-digested plasmid using T4 DNA ligase (NEB). Four microliters of the ligation mixture were transformed into XL Gold-competent *E. coli*. Correct integration of the gRNA cassette was verified by colony PCR employing the reverse oligonucleotide in combination with a forward primer upstream of the plasmid.

The DCX homology-directed repair template (PbDCX_PbU6_3HA_gRNA) was commercially synthesised (Genscript). The full sequence of PbDCX_PbU6_3HA_gRNA is shown in **Data S1C** and **D**. For tagging constructs, 600 bp homology arms flanking the gRNA cleavage site were incorporated, positioning the break near the 3′ end of the coding sequence, where a triple HA tag was inserted before the stop codon. The region harbouring the gRNA recognition site was recodonized to prevent re-cleavage of the modified locus. The DCX::HA fragment was ligated into the plasmid containing the gRNA using HindIII and NcoI (NEB). The ligation products were introduced into XL Gold *E. coli*, and transformants were subsequently screened for the presence of the insert by colony PCR. The complete plasmid was then sequenced (with PlasmidSaurus).

### Transfection of DCX CRISPR/Cas9 vector in *P. berghei*

Schizonts of *Plasmodium berghei* ANKA 2.34 were purified on a 55% Histodenz gradient and electroporated with 10 µg of plasmid DNA (one sgRNA construct) using the Amaxa 4D-Nucleofector (program FI-115; Lonza) in P3 Primary Cell Nucleofector Solution with Supplement 1 (Lonza). Transfected parasites were allowed to reinvade erythrocytes for 1 h prior to intraperitoneal injection into recipient mice. Selection was applied 24 h post-transfection by administering pyrimethamine (0.07 mg/ml; Sigma) in drinking water for five days. Clonal parasite lines were obtained by limiting dilution (1–2 parasites per mouse, 5–10 mice) and verified by PCR to confirm correct integration and exclude wild-type alleles. All experiments were performed in female CD1 outbred mice (6–12 weeks old) obtained from Charles River Laboratories. For transfections, erythrocytes used for parasite reinvasion were obtained from donor mice pretreated with phenylhydrazine three days before collection to induce reticulocytosis.

### P. falciparum growth assays

The growth of *PfDCX-HA:cKO* parasites was assessed as previously described^35^. Briefly, parasites were synchronised as early rings (2-3h post-invasion) and treated with 0.1% DMSO (vehicle) or 100nM rapamycin (LC Laboratories) all along. Parasitaemia was monitored microscopically during 3 replication cycles using Haemacolour-stained thin blood films at day 0, 2, 4 and 6. Parasitaemia was calculated at each time point by randomly selecting 10 to 12 microscopic fields and calculating the number of infected RBCs/total number of RBCs × 100.

### *P. berghei* ookinete culture and purification

Gametocyte-infected blood was collected from *Plasmodium berghei* PbDCX-HA-infected mice and cultured in ookinete medium consisting of RPMI 1640 (HEPES, L-glutamine, without bicarbonate; 0.164 g/10 ml), hypoxanthine (0.5 mg/10 ml), sodium bicarbonate (20 mg/10 ml), penicillin–streptomycin (100 µl/10 ml; 5,000 U/ml stock), and xanthurenic acid (100 µM final concentration), adjusted to pH 7.6 and sterile-filtered. Prior to use, the medium was supplemented with 20% (v/v) fetal bovine serum. Cultures were maintained at 19°C for 20–24 h to allow ookinete development.

Mature ookinetes were purified by centrifugation over a 17% (w/v) Histodenz cushion, while zygotes and stage III ookinetes were purified using a 14% (w/v) Histodenz cushion. Infected blood–culture mixtures were layered onto the gradient and centrifuged at 1,360 × g for 10 min at room temperature without brake (3/3). The ookinete-enriched fraction at the interface was collected, washed once in ookinete medium, and resuspended for downstream applications.

### Immunofluorescence assay of PfDCX-HA schizonts

Schizonts were collected using standard Percoll enrichment as described above, then fixed in 4% paraformaldehyde (PFA) and 0.0065% glutaraldehyde at room temperature for 20 minutes. PBS-resuspended samples were adhered to PolyD-coated coverslips. Cells were permeabilised with Triton X-100 for 5 minutes and washed with PBS1x. After blocking with 2%BSA for 30 minutes, primary antibodies were incubated overnight at 4°C (rat anti-HA 1:1000; guinea pig anti-α-Tubulin and guinea pig anti-β-Tubulin 1:2000). After 3 washes with PBS1x, secondary antibodies were added for 3 hours, at room temperature, in the dark (anti-rat 488 1:1000, anti-guinea pig 647 1/1000). Following 3 washes with PBS1x, slides were mounted with antifade agent (Vectashield) supplemented with the fluorescent DNA stain 4′,6-diamidino-2-phenylindole (DAPI, 1 μg/ml) (Vector Laboratories).

### *P. falciparum* induction of gametocytogenesis and gametocyte culture

The role of PfDCX in *in vivo* transmission was assessed by inducing gametocytes as described previously^36^. Briefly, gametocyte cultures were initiated from 2% asexual blood-stage cultures at a haematocrit of 4%. Gametocytes were cultured in gametocyte medium (RPMI-1640, 2.78 g/L sodium bicarbonate, 4 g/L glucose, 5.9 g/L HEPES, with 5% human AB+ serum, 0.5% Albumax, 3.7% 100xHT supplement (ThermoFisher), 30 mg/ml L-glutamine) at 37°C, which was replaced daily for 14 days. On day 14, exflagellation was triggered (as described below) to confirm gametocyte maturity. Cultures showing at least 0.2% exflagellation (of total cells) were used for standard membrane feeding assays.

To study the role of PfDCX *in vitro*, highly synchronised gametocytes were induced as previously performed^35^. Briefly, asexual parasites were cultured in Albumax-complemented medium and synchronised over two replication cycles using 5% sorbitol, as described above. When reaching an old ring stage, parasites were treated with a minimal fatty acid medium containing RPMI (Gibco, lot 2436708) complemented with bovine serum albumin (BSA; 390 mg/100 ml), 30 μM oleic acid, and 30 μM palmitic acid. Twenty-four hours post-induction, when parasites were schizonts/young rings, serum-complemented medium containing RPMI (Gibco, lot 2436708), hypoxanthine (200 µM; Sigma), 10% human AB serum, and gentamycin (40 µg/ml; Gibco) was used. Parasites were treated with rapamycin (100nM) or 0.1% DMSO for 48 hours. One day later, referred to as gametocyte day 1 (GamDay1), early gametocytes were fed with serum/N-acetylglucosamine (GlucNac, Sigma A3286-100G) medium (serum medium supplemented with 50 mM GlucNac) for 7 consecutive days to eliminate asexual parasites^37^. From GamDay6 onwards, parasites were cultured on a heating plate to maintain 37°C. Parasites were fed with serum medium from day 8 and kept until day 13 when they reached full maturity (gametocyte stage V).

### *P. falciparum* exflagellation assays

Exflagellation assays were performed as described previously^35^. Briefly, gametocytaemia was determined by making thin blood smears of gametocyte cultures at GamDay 13, (when reaching full maturity). 500µL of resuspended cultures were taken, spun down at 300g for 30 seconds. 2 µL of the pellets were taken and resuspended in 50 μl of 1× exflagellation medium (serum medium containing 100 μM XA). The activation mixes were incubated in a Neubauer chamber in the dark and in a humid chamber for 15 minutes at room temperature. The percentage of exflagellating gametocytes was calculated as follows:

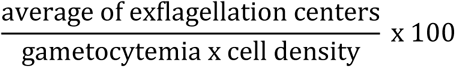

### Induction of developing *P. falciparum* ookinetes *in vitro*

Retorts were obtained *in vitro* as previously described^22^, with minor modifications. *P. falciparum* stage V gametocyte cultures exhibiting acceptable exflagellation (>0.2%) were pelleted at 500 × g for 5 min at 37 °C. The supernatant was removed, and the pellet was resuspended in an equal volume of human serum at room temperature, resulting in a final haematocrit of 50%. The suspension was gently mixed by inversion and incubated for 2 h at room temperature to allow fertilisation of activated male and female gametes. Subsequently, complete gametocyte culture medium (RPMI-1640 supplemented with 25 mM HEPES, 2 mM L-glutamine, 2 g/L sodium bicarbonate, 50 mg/L hypoxanthine; pH 8.2) was added at 10× the pellet volume. After 26 h of incubation at room temperature, developing ookinetes displaying an apical protrusion were visualised by live imaging using an anti-Pfs25-Cy3 antibody (1:500)^38^.

### Immunofluorescence assay of PfDCX-HA retorts

Samples were fixed with a final concentration of 4% PFA by adding an equal volume of 8% PFA in PBS to retort cultures for 20 minutes at room temperature. 100 μl of the fixed cell suspensions was added to wells of a 24-well plate containing 1ml PBS and a poly-l-lysine-coated glass coverslip, and the plate was incubated overnight at 4°C, allowing cells to adhere. Coverslips were placed on a hydrophobic Parafilm surface for immunofluorescence staining in a humidified, dark chamber. Cells were permeabilised with 0.1% Triton X-100 in PBS for 10 minutes and washed three times in PBS. Coverslips were incubated in blocking buffer (PBS with 10% fetal bovine serum, FBS) for 1 hour. After washing, cells were labelled with anti-Pfs25-Cy3^39^, 11,000 dilution) and anti-HA antibody (3F10, 1:250 dilution) for 1 hour. Cells were then wash three times in PBS and then stained with anti-rat Alexa 488 (1:1,000 dilution) before a final 3 washes. Coverslips were mounted onto glass slides using Vectashield with DAPI and sealed with nail polish. Cells were visualised by epifluorescence microscopy using a Nikon TiE2 widefield microscope at x100 objective with x1.5 zoom lens. Z-stack images were captured and processed by deconvolution using NIS Elements software.

### Phenotypic analysis of PfDCX-KO developing *in vitro* ookinetes

*PfDCX-HA:cKO* gametocytes treated with DMSO or 100nM rapamycin were induced to form *in vitro* developing ookinetes as described above. Parasites were stained with anti-Pfs25-Cy3 antibody (1:500) and visualised by live imaging using the Nikon TiE widefield microscope. Developing ookinetes were classified morphologically as thick (short, curved apical protrusion) or thin (elongated, slender protrusion). Three independent biological replicates were performed, with 1,450–2,722 cells counted per replicate. The percentage of developing ookinetes was calculated as follows:

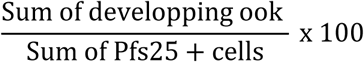

The percentage of thick or thin developing ookinetes was calculated as:

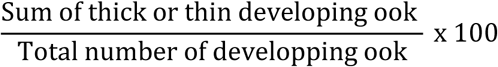

### Standard Membrane Feeding Assays

*PfDCX-HA:cKO* gametocyte cultures treated with DMSO or 100 nM rapamycin and exhibiting adequate exflagellation (>0.2%) were pelleted by centrifugation. Pellets were resuspended in an equal volume of human serum at 37 °C, yielding a final haematocrit of 50% to generate a blood meal. Standard membrane feeding assays (SMFAs) were performed as previously described^40^. Briefly, gametocytes were fed to *Anopheles coluzzii* mosquitoes for 30 min at 37 °C through a Parafilm membrane. Engorged mosquitoes were maintained at 28 °C with 60–80% relative humidity. Midgut dissections were performed at day 7 post-infection, and the number of oocysts per mosquito was determined using a brightfield microscope (objective 10X). Two independent replicates were conducted, with 25–58 mosquitoes infected per replicate. Oocyst prevalence was calculated as the proportion of infected mosquitoes:

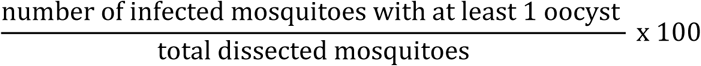

Infection with *PfNF54* parasites reached a prevalence of 76.6%, confirming that the required procedural standard was met (**Fig. S2**).

### Quantitative analysis of stage V gametocyte morphology

Stage V gametocyte cultures were prepared as described above. Haemacolor-stained thin blood smears were used to determine gametocytaemia and imaged with a Zeiss Axio Observer Z1 microscope equipped with a 63× oil objective. Four biological replicates of PfDCX-HA and three of PfDCX-KO were analysed, with 200 gametocytes quantified per sample using QuPath. A machine-learning algorithm was trained to identify infected RBCs containing gametocytes based on Haemacolor staining. Following gametocyte detection, morphology was assessed by calculating the perimeter-to-circularity ratio.

### Sample preparation for U-ExM

*P. falciparum* schizonts and gametocytes (stage IV and stage V) were collected using a 63% Percoll gradient (GE Healthcare, Life Sciences). For gametocytes, the collection process was done at 37°C to avoid premature activation.

After purification, schizonts were resuspended in 50-100µL incomplete RPMI (Gibco, lot 2436708), and sedimented into 12mm round Poly-D-Lysine (A3890401, Gibco) coated coverslips for 15 minutes. Coversplits were submerged in 100% methanol at -20°C for 7 minutes.

*P. falciparum* gametocytes were either directly fixed (stage IV and non-activated stage V gametocytes) or were activated using a human serum-complemented medium (10% human serum, gentamicin (40 µg/ml; Gibco), hypoxanthine (200 µM; Sigma)) containing 100µM Xanthurenic acid for 15 minutes under rotation at room temperature and subsequently fixed in 2% formaldehyde/ 0.0065% glutaraldehyde for 10 minutes. Gametocyte samples were then sedimented into 12mm round Poly-D-Lysine (A3890401, Gibco) coated coverslips for 15 minutes. Samples were subsequently used for U-ExM.

*P. falciparum in vivo* ookinetes were purified from dissected mosquito midguts, which were smashed and layered into a 14.1% (w/v) Gentodenz® (Gentaur) gradient (20mins, 800g, at room temperature). A thin layer of cells was recovered, washed in incomplete RPMI and settled in PolyD-coated coverslips for 30 minutes. Coverslips were fixed in -20°C methanol for 7 minutes. *P. berghei* ookinetes were sedimented on PolyD-lysine-coated coverslips (A3890401, Gibco) at room temperature for 10 minutes and fixed for 6 minutes with 2% methanol-free formaldehyde (Thermo Fisher Scienfitix, 28906) in PBS1X, followed by a fixation for 1 minute in methanol at -20°C. Cells were then washed 3 times with 100mM glycine in PBS.

### U-ExM

After being settled in PolyD-coated coversplits, samples were used for U-ExM as previously^28^. The subsequent steps were carried out as follows:

1. **Protein anchoring**: Coverslips were incubated for 5 h at 37 °C in a solution of 1.4% formaldehyde (FA) and 2% acrylamide (AA).
2. **Gelation**: Samples were polymerised for 1 h at 37 °C in a monomer solution (23% sodium acrylate, 10% AA, 0.1% BIS-AA in PBS) supplemented with 10% ammonium persulfate (APS) and 10% TEMED.
3. **Denaturation:** Gels were incubated at 95 °C for 1 h 30 min.
4. **Gel expansion and antibody labelling**: Following denaturation, gels were incubated in ddH₂O at room temperature for 30 min and then left overnight in ddH₂O for complete expansion. The next day, gels were washed twice in PBS1X (15 min each) to remove excess water, incubated with primary antibodies for 3 h at 37 °C with shaking (160 rpm), and washed three times for 10 min in PBS-Tween 0.1%. Secondary antibody incubation was then performed under the same conditions, followed by three additional washes. Immediately after antibody staining, gels were incubated with 594 NHS-ester (Merck, 08741; 10 μg/mL in PBS1X) for 1 h 30 min at room temperature with shaking, washed three times in PBS-Tween 0.1% (15 min each), and expanded a second time overnight in ddH₂O before imaging.
5. **Sample mounting and imaging:** Gel pieces (∼1 cm × 1 cm) were cut and mounted on 24 mm round Poly-D-Lysine–coated coverslips (Gibco, A3890401) to prevent gel drift. Coverslips were then fitted into metallic O-ring imaging chambers (Okolab, RA-35-18 2000–06) and imaged.

Images were acquired on a Leica TCS SP8 microscope with an HC PL Apo 100x/1.40 oil-immersion objective. The lightning mode was used to generate deconvolved images. Z-stacks were captured between frames using HyD as the detector. Images were processed using ImageJ, LAS X, and Imaris 9.6. Antibodies, stains and reagents used for U-ExM are listed in **Table S2** and **Table S3**, respectively.

### Iterative U-ExM

*P. berghei* ookinetes were purified, settled in PolyD-coated coverslips and fixed as described above. Coverslips were processed for iU-ExM as previously described^23^. Briefly, 12-mm coverslips were incubated overnight at 37 °C in anchoring solution (1.4% formaldehyde, 2% acrylamide in 1× PBS). Coverslips were then mounted onto a glass slide with glue, and a 16-mm spacer (Sunjin Lab) was placed around the sample to create a chamber. Gelation solution (10% acrylamide, 19% sodium acrylate (SA), 0.1% DHEBA, 0.25% tetramethylethylenediamine (TEMED)/ammonium persulfate (APS)) was added to the chamber, which was sealed with a 24-mm coverslip. Gelation was carried out for 15 min on ice followed by 45 min at 37 °C.

After polymerisation, the 24-mm coverslip and spacer were removed, and the gel-embedded coverslip was gently detached from the slide and incubated in 2 mL denaturation buffer (200 mM SDS, 200 mM NaCl, 50 mM Tris base, pH 6.8) in a 6-well plate under shaking to release the gel from the coverslip. The gel was then transferred to an Eppendorf tube containing denaturation buffer and incubated at 85 °C for 1 h 30 min. Following denaturation, gels were placed in a 10-cm dish and washed in three consecutive ddH₂O baths of 30 min each.

A 1.5 × 1.5 cm gel punch was excised and incubated in 1× PBS to shrink the gel prior to immunostaining. Primary antibodies were incubated overnight at 4 °C in PBS containing 2% BSA under gentle shaking, followed by three washes with PBS containing 0.1% Tween-20 (5 min each). Secondary antibodies were then incubated for 2 h 30 min at 37 °C in PBS/2% BSA under shaking, followed by three additional washes with PBS/0.1% Tween-20 (5 min each). NHS-ester was subsequently incubated for 1 h at 37 °C in 1× PBS under shaking, followed by three washes with PBS/0.1% Tween-20 (5 min each).

The stained gel punch was transferred to a 12-well plate on ice and incubated for 25 min with 2 mL of activated neutral gel solution (10% acrylamide, 0.05% DHEBA, 0.1% APS/TEMED in ddH₂O) with the lid open under shaking. The gel was then placed onto a microscope slide, excess monomer solution was gently removed using tissue paper, and the gel was covered with a 22 × 22 mm coverslip. Polymerisation was carried out in a humid chamber for 1 h at 37 °C.

Following polymerisation, the coverslip was removed and the gel was transferred to a new 12-well plate for a second anchoring step using anchoring solution (1.4% formaldehyde, 2% acrylamide in 1× PBS) for 3 h at 37 °C. The gel was then incubated on ice under shaking for 25 min in 2 mL of second expansion monomer solution (10% acrylamide, 19% SA, 0.1% BIS, 0.1% TEMED/APS). The gel was transferred onto a glass slide, excess monomer solution was removed, and the gel was covered with a 22 × 22 mm coverslip and incubated in a humid chamber for 1 h at 37 °C.

After polymerisation, the gel was incubated in 200 mM NaOH for 1 h at room temperature under shaking, followed by two washes in 1× PBS (15 min each). Finally, the gel was expanded by incubation in three successive ddH₂O baths of 2 h each. Antibodies and stains used are listed in **Table S2**. Reagents used were previously described^23^.

### High-throughput immunofluorescence assay in activated gametocytes

Gametogenesis was assessed in PfDCX-HA and PfDCX-KO activated gametes as described previously^25^. Briefly, 150µl of gametocyte cultures were taken and incubated with the same volume of 1× exflagellation medium (serum medium containing 100 μM XA), for 20 minutes at room temperature. Samples were fixed with a final concentration of 4% PFA by adding an equal volume of 8% PFA, for 20 minutes at room temperature. Then, 100 μl of the fixed cell suspensions were added to wells of a 24 well plate containing 1ml PBS and a poly-l-lysine-coated glass coverslip and incubated overnight at 4°C, allowing cells to adhere. Coverslips were placed on a hydrophobic Parafilm surface for immunofluorescence staining in a humidified, dark chamber. Cells were permeabilized with 0.1% Triton X-100 in PBS for 10 min and washed three times in PBS. Coverslips were incubated in blocking buffer (PBS with 10% fetal bovine serum, FBS) for 1 h. Microgametes were labelled with an anti–α-tubulin antibody (clone DM1A, Sigma; raised against chicken embryo brain tubulin but cross-reactive with multiple species) diluted 1:500 in blocking buffer for 1 h, followed by washes and incubation with an anti-mouse Alexa Fluor 488 secondary antibody (Thermo Fisher) diluted 1:1,000 in blocking buffer for one hour. After final washes, coverslips were mounted onto glass slides with VectaShield containing DAPI 1 μg/ml (Vector Laboratories) to visualise DNA.

The Nikon Tie2 fluorescence microscope at an x100 oil objective and 1.5x zoom was used to image the slides. For each sample, tubulin-positive, DAPI-positive stained gametes were imaged randomly. For each field of view images were acquired with a Z-stack at 0.3 μm intervals and subsequently deconvolved and processed into maximum intensity projections using Nikon NIS Elements software. In the raw data, cells were initially detected using an automated ICY Bioimage Analysis pipeline based on tubulin signal intensity, which defined a region of interest (ROI) around each cell. All ROIs were then reviewed and adjusted manually when necessary. Each raw image yielded between 1 and 8 analysable cells. Analysis was done using the PhIDDLI pipeline as described in^25^.

PhIDDLI, a pre-trained machine learning image analysis pipeline was implemented in Python 3.8 and executed via command line on a 64-bit Ubuntu virtual machine running in Oracle VM VirtualBox 6.1. Output data were visualized as an interactive webpage in Mozilla Firefox and exported as comma-separated values (CSV) files for downstream analysis.

### Sex ratio determination of *P. falciparum* gametocytes by immunofluorescence assays

Sex ratio of *PfDCX-HA*:cKO/DiCre parasites treated with or without rapamycin was determined as previously described^35^. Briefly, 500 μl of gametocyte cultures (Gamday10) were spun down (400g, 1 min) and used for making thin blood smears. Slides were fixed with 60% methanol/40% acetone (at −20°C) for 3 min and stored at −80°C. Slides were used for immunofluorescence assays (IFAs). Cells were hydrated with PBS1X and blocked with 3% BSA in PBS 1× for 1 hour. Primary antibodies were added (1:1000; rabbit anti-Pfg377)^41^ and incubated for 1 hour, following 6 washes with PBS1X. Secondary antibodies (1:250; anti-rabbit 568) were incubated for 1 hour in the dark. After washing with PBS1x, the antifade agent (Vectashield) supplemented with the fluorescent DNA stain 4′,6-diamidino-2-phenylindole (DAPI) (1 μg/ml) was used, and slides were sealed. Image acquisition was done with the LSM800 Airyscan using the 40x objective. A minimum of 100 DAPI-positive cells were counted per condition.

### Transmission electron microscopy (TEM) of *P. falciparum* gametocytes

Stage IV and stage V gametocytes were collected with a 63% Percoll gradient (GE Healthcare, Life Sciences). Stage V gametocytes were activated with 1× exflagellation medium (serum medium containing 100 μM XA). Samples were fixed with a mixture of 2.5% glutaraldehyde and 2% paraformaldehyde for 1 hour at room temperature. After fixation, samples were washed three times in PBS (1×) and stored in PBS (1×) at 4 °C until further use. Pellets were washed three times for 5 min each in 0.1 M sodium cacodylate buffer (pH 7.4), followed by post-fixation in 1% osmium tetroxide (Electron Microscopy Sciences) with 1.5% potassium ferrocyanide in 0.1 M sodium cacodylate buffer (pH 7.4) for 1 h. Samples were then incubated in 1% osmium tetroxide alone in 0.1 M sodium cacodylate buffer (pH 7.4) for 1 h, washed twice in double-distilled water (5 min each), and, to reduce sample loss in subsequent processing steps, cell pellets were embedded in 2% agar. After solidification, small (∼1 mm³) cubes were cut out and samples were *en bloc* stained with 1% uranyl acetate (Electron Microscopy Sciences) in ddH₂O overnight at 4 °C, washed twice in ddH₂O (5 min each), and dehydrated through a graded ethanol series (2 × 50%, 1 × 70%, 1 × 90%, 1 × 95%, 2 × 100%; 10 min each). Samples were then washed once in propylene oxide for 10 min. Dehydrated samples were infiltrated with 50% EPON epoxy resin diluted with propylene oxide for 1 h, followed by 100% fresh epoxy resin overnight. Finally, samples were embedded in pure fresh epoxy resin and placed in gelatin capsules, then polymerised in an oven at 65 °C for 24–48 h.

Using an ultramicrotome (Leica UC7, Austria) and a diamond knife (DiATOME, Switzerland), 50 nm-thick ultrathin sections were cut and collected on copper 100 mesh hexagonal EM grids coated with Formvar plastic support film and carbon. Sections were examined with Tecnai 12 G2 TEM (FEI, Netherlands) equipped with a side-mounted MegaView III CCD camera (Olympus-Soft Imaging Solutions) and Morgagni TEM (FEI, Netherlands) equipped with a side-mounted Veleta CCD camera (Olympus-Soft Imaging Solutions). Both microscopes were operated at an acceleration voltage of 80 kV, and both cameras were controlled by iTEM acquisition software (Olympus-Soft Imaging Solutions). All TEM sample preparation and imaging were performed by the Electron Microscopy Facility (PFMU) at the Medical Faculty, University of Geneva, Switzerland. The distance between SPMTs was quantified by counting the number of microtubule rafts within a defined length across a cell traversal cross-section.

### Quantification of the polyglutamylation and α/β-Tubulin signals by corrected total cell fluorescence (CTCF) in macro- and microgametocytes

Image analysis was performed using the Fiji software. Maximum intensity projections were generated for each acquired channel. Brightness and contrast were adjusted identically across all samples for each channel. A threshold based on the NHS-ester channel was manually applied to delineate individual cells. For each cell, the integrated density, area and mean background were measured. The corrected total cell fluorescence (CTCF) was calculated as:

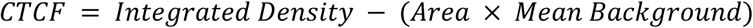

The “integrated density” is the sum of all pixel intensities within the selected cell area, while “Area × Mean Background” represents the estimated contribution of background fluorescence within that same region. Mean background intensity was measured in a cell-free region of the same image. Statistical analysis was performed using a two-way ANOVA in GraphPad Prism.

### Purification of stage IV gametocytes for Cryo-Electron tomography

As for TEM, stage IV gametocytes were collected with a 63% Percoll gradient (GE Healthcare, Life Sciences) and kept at 37°C in incomplete RPMI (Gibco, lot 2436708).

### Preparation of Gametocytes for Cryo-Electron Tomography

Gametocytes were kept at 37 °C at all times through use of a heat block. Quantifoil R 1/4 grids (Quantifoil Micro Tools) were plasma cleaned using a GlowQube (Quorum Technologies) on both sides using residual air prior to freezing. Plunge freezing was carried out using a Leica GP2 plunge freezer (Leica Microsystems) with the chamber set to 70% humidity and 37 °C. 3 μl of gametocyte solution was added to the front side of the grids, and 1 μl of gametocyte medium was added to the back side to aid in the flow-through of material and ensure more even and reproducible blotting. Grids were blotted from the back for 3 seconds before immediately proceeding with plunging into liquid ethane at -185 °C.

Prior to Focused Ion Beam (FIB) milling, grids were clipped into FIB-specialised Autogrids (Thermo-Fisher Scientific). FIB milling was carried out using an Aquilos 2 system (Thermo-Fisher Scientific). GIS deposition of 90 s and platinum sputtering at 10 pA and 1 kV for 30 s were performed inside the instrument before milling. Micro-expansion joints^42^ were milled to minimise lamella bending. Milling was carried out at 10 degrees in a 45-degree shuttle. Rough milling was carried out iteratively at currents of 1 nA, 500 pA, and 300 pA, dropping the current as the lamella approached its final thickness. Polishing down to ∼200 nm was carried out using 100 and 50 pA currents. After milling all sites on a grid, another round of sputtering at 1 kV and 10pA for 5 s was performed.

### Cryo-Electron Tomography Tilt Series Collection

Data collection was carried out on a Titan Krios (Thermo-Fischer Scientific), equipped with a Falcon 4i Camera (Thermo-Fischer Scientific) and a SelectrisX energy filter (Thermo-Fischer Scientific) set at zero loss. Tomo5 software (Thermo-Fischer Scientific) was used for microscope operations. Tilt series collection was carried out using a dose symmetric scheme^43^ starting with a -10 degree pretilt, acquiring images every 2 degrees between -60 and +40 degrees. Image acquisition was at 3 Å/px with a target defocus of -7 μm for a final electron dose of 120 e-/Å2. 32 tilt series were successfully collected in this manner.

### Tomogram reconstruction

Frames were preprocessed with WarpTools^44^ by motion correction, defocus and CTF estimation. The tilt series was aligned using IMOD’s^45^ tilt alignment program, followed by tomogram reconstruction in WarpTools along with odd-even frames for denoising. The tomograms were further denoised using cryo-CARE^46^ for visualisation.

### Microtubule subtomogram averaging (STA)

The microtubules were manually traced by tracing a line through their centres in IMOD. STA was performed in Dynamo^47^ using the dynamoMT scripts (https://github.com/builab/dynamoMT) along with a set of homemade scripts. First, the subtomograms were picked every 8 nm along the length of the microtubules. Then, subtomograms belonging to the same microtubule were roughly aligned and averaged. Each subtomogram was then assigned a randomised rotation angle to reduce the effects of the missing wedge. Subsequently, alignment was performed for per-microtubule averages with updated symmetry parameters, followed by classification based on their protofilament numbers using custom scripts (13-18).

### Electrophoresis and immunoblotting

Samples were saponin-lysed (10 mins, 4°C) and centrifuged for 5 mins at full speed at 4°C. Pellets were lysed in RIPA buffer containing: 150mM NaCl, 5mM EDTA, pH 8, 50mM Tris, pH 7.5, 1% NP40, 0.5% sodium deoxycholate, 0.1% SDS. Protein separation was done on Novex^TM^ Tris-Glycine gel 10-20% (XP10205BOX, Invitrogen) and transferred to nitrocellulose membranes by electroblotting (Amersham Protran premium 0.45µm membrane NC, 10600003). Blots were blocked for 1 h at room temperature with 5% milk in PBS-Tween 0.05% and primary antibodies added overnight at 4°C. After 3 washes with PBS-Tween 0.05%, secondary antibodies were added for 1 hour at room temperature, and washed as previously. Membranes were revealed with the ECL Prime Western Blotting Detection Reagents (RNP2232, Cytiva). Blots were subsequently incubated with anti-actin antibody as a loading control. Antibodies used are listed in **Table S2**. Source data for western blot analyses are provided as Supplementary **Data S2** and **S3**.

### Ethics statement

The animal work performed in this study has undergone ethical review and was approved by the Swiss Federal Veterinary Office under authorisation number GE194. Six- to ten-week-old mice were obtained from Charles River Laboratories (France), and females were used for all experiments. Mice were specific pathogen-free (including *Mycoplasma pulmonis*) and subjected to regular pathogen monitoring by sentinel screening. They were housed in individually ventilated cages furnished with a cardboard mouse house and Nestlet, maintained at 21 ± 2°C under a 12-hour light/dark cycle, and given commercially prepared autoclaved dry rodent diet and water *ad libitum*.

### Statistical analyses

Statistical analyses were performed using GraphPad Prism version 10.5. Differences between the two groups were assessed using an unpaired two-tailed Student’s t-test or a paired t-test when comparing parasite lines treated in parallel with a drug and the vehicle control. For comparisons involving more than two groups, one-way ANOVA followed by either Tukey’s pairwise comparison test or Šídák’s multiple comparisons test was applied, as indicated for each experiment. When comparing categorical distributions across groups, a χ² test or a Fisher’s exact test was used. P-values are reported, with statistical significance defined as p < 0.05. All experiments were performed in two to three independent biological replicates, as indicated in the figure legends. Representative images from one replicate are shown.

## Acknowledgments.

We thank Dr Judith Green (The Francis Crick Institute, London, UK) for sharing the *P. falciparum* MyoB-GFP line, as well as Dr Eilidh Carrington from Prof Till Voss laboratory (SwissTPH, Switzerland) for sharing her transfection protocol. We further thank Natacha Klages Jemelin for technical support and Dr Konstantinos Kousis (LSHTM, UK) for support with image acquisition of expanded *P. falciparum* ookinetes. We thank the excellent support of the core facilities from the Faculty of Medicine of the University of Geneva: Dr François Prodon and Olivier Brun (Bioimaging Facility); Dr Bohumil Maco and Pilar Ruga (Pôle Facultaire de Microscopie Ultrastructurale). We also thank Lindsay Stewart and Mojca Kristan for performing the standard membrane feeding assays (Human Malaria Transmission Facility, LSHTM, UK). This work was supported by the Swiss National Foundation (grants 310030_208151 and CRSII5_198545 to MB), the Fondation privée des Hopitaux Universitaires de Genève (grant RC08-16 to MB, VH, and PG), the ERC Consolidator Grant ISAC, administered in Switzerland by SERI (contract MB22.00075 to PG), a UKRI Medical Research Council Career Development Award (MR/V010034/1 to MJD), a Medicines for Malaria Venture award (RD-21-1003 to MJD), a BBSRC London Interdisciplinary Biosciences Programme Studentship to RB, and by a grant from the BRYN TURNER-SAMUELS Foundation, with support from the Foundation for Research in Biology and Medicine (support for GenExM to VH, PG and OM).

