## Supplementary material for "DCX enables branching of subpellicular microtubules in *Plasmodium falciparum* gametocytes and is required for mosquito colonisation"

Emma Ganga *et al.*

**This file includes:**

**Supplementary Figures: S1 to S5**

**Supplementary Tables: S1 to S3**

**Supplementary Data: S1 to S3**

### Supplementary figures

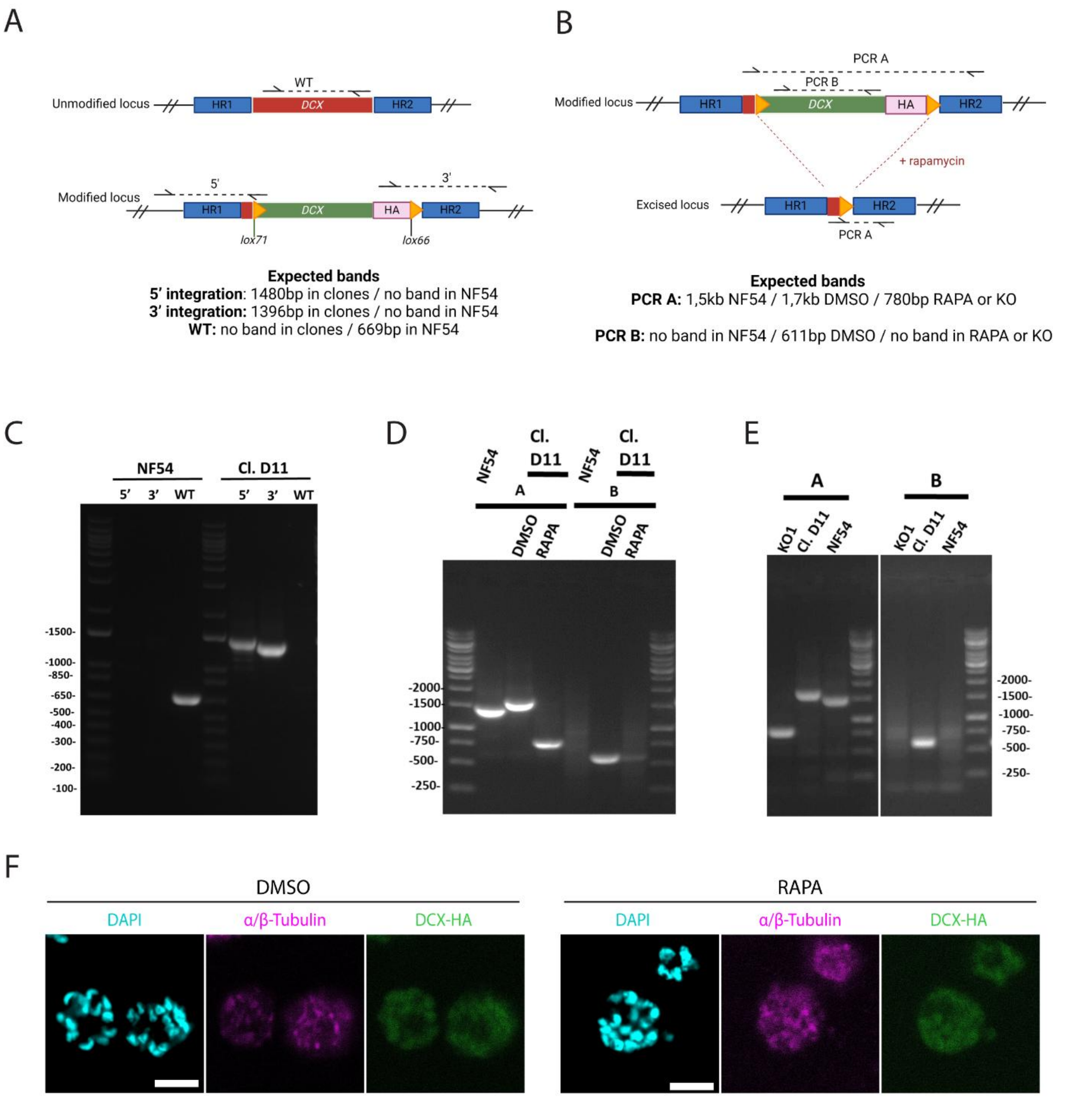


**Figure S1. Generation and validation of the PfDCX-HA:cKO line and rapamycin-induced excision of *PfDCX-HA*. (A)** Schematic of the PfDCX-HA:cKO targeting strategy in *P. falciparum*. The endogenous *PfDCX* locus (red) is truncated and replaced by a recodonised *PfDCX* gene (green) flanked by two lox sites (lox71 and lox66) and is tagged by 6 HA epitopes. Correct integration was assessed by PCR using primer pairs amplifying the wild-type (WT), the 5' integration, and the 3' integration. Expected amplicon sizes are 1480 bp (5' integration) and 1396 bp (3' integration). No WT band is expected in transfectant parasites, whereas a 669 bp band is expected in the parental NF54Dicre line. HR, homology region. **(B)** PCR strategy to assess rapamycin-induced excision of PfDCX-HA. PCR A amplifies the recodonised *DCX* locus and flanking homology regions, yielding bands of 1.5 kb in the parental NF54DiCre line, 1.7 kb in DMSO-treated parasites, and 780 bp following rapamycin-induced excision. PCR B amplifies the recodonised *DCX* sequence, producing a 611 bp band in DMSO-treated parasites, with no band expected after rapamycin treatment or in the parental line. **(C-E)** Agarose gel electrophoresis confirming correct PCR products. **(C)** Correct 5' and 3' integration in the PfDCX-HA:cKO clone D11 (Cl. D11), which was used for all experiments in this study. **(D)** Correct excision of *PfDCX-HA* in PfDCX-HA:cKO Cl. D11 following rapamycin treatment, and **(E)** in the PfDCX-KO clone 1 (KO1). **(F)** Immunofluorescence images of PfDCX-HA:cKO schizonts treated with or without rapamycin and stained with DAPI (DNA), α/β-tubulin, and DCX-HA. Scale bar, 4μm.

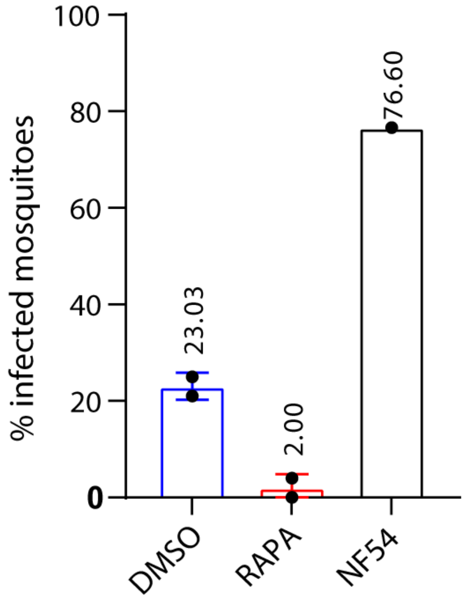


**Figure S2. Oocyst prevalence in mosquito midguts infected with NF54 gametocytes or PfDCX-HA:cKO gametocytes treated with DMSO or rapamycin.** NF54 parasite infection showed a 76.6% prevalence, confirming that the infection met the required procedural standard (n = 25 to 58 mosquitoes per condition per replicate).

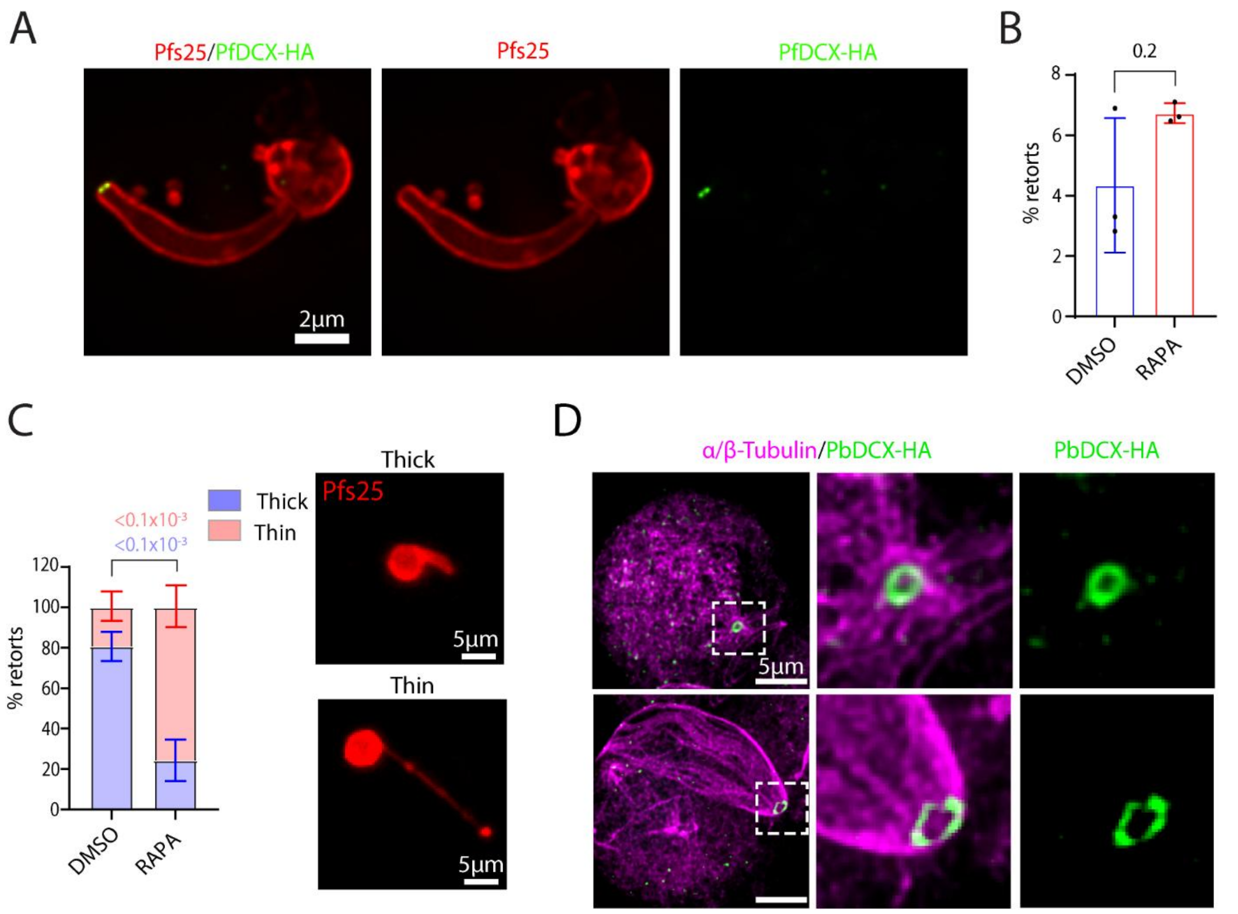


**Figure S3. PfDCX localises to the apical end of *in vitro* retorts and affects retort morphology. (A)** Representative immunofluorescence image of an *in vitro P. falciparum* retort expressing the surface membrane marker Pfs25 and PfDCX-HA, which localises to the apical end. Scale bar: 2µm. **(B)** Proportion of retorts formed *in vitro* from gametocyte cultures treated with DMSO or rapamycin (mean ± SD, n=3 biological replicates, paired two-tailed *t* test). **(C)** Proportion of *in vitro*-formed retorts from DMSO- or rapamycin-treated gametocyte cultures displaying either a thicker curved protrusion or a thin, elongated protrusion (mean ± SD, n=3 biological replicates, Fisher's exact test, two-sided, p-values <0.0001). Representative images of retorts stained with the Pfs25 membrane marker illustrate typical and thin, elongated protrusion morphologies. Scale bar: 5µm. **(D)** Representative U-ExM images of a *P. berghei* zygote (top) and a developing stage III ookinete (bottom) expressing PbDCX-HA at the apical end of SPMTs (α/β-tubulin). Scale bar: 5µm.

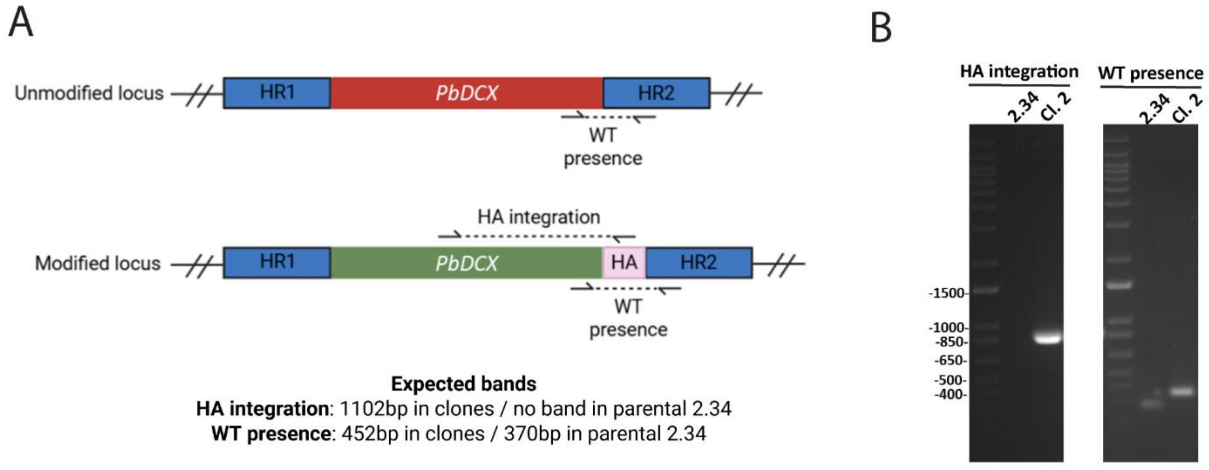


**Figure S4. Generation and validation of the PbDCX-HA line. (A)** Schematic of the targeting strategy used to generate PbDCX-HA in *P. berghei*. The endogenous *PbDCX* locus (red) is replaced by a recodonised PbDCX gene (green) containing 3HA epitopes. PCR assessed correct integration: the HA integration amplicon yields a band of 1102 bp in the PbDCX-HA clone 2 (Cl. 2), and no band in the parental line 2.34, while the WT presence amplicon produces bands of 452 bp in PbDCX-HA Cl. 2 and 370 bp in the parental line. **(B)** Agarose gel electrophoresis confirming the expected PCR band sizes.

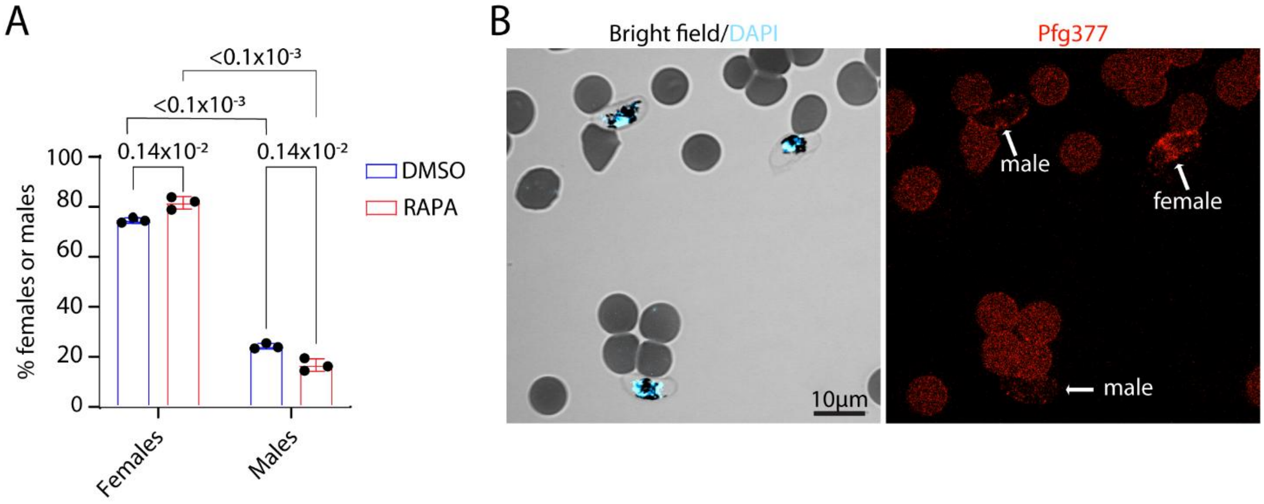


**Figure S5. *PfDCX* deletion alters the sex ratio. (A)** Sex ratio analysis of PfDCX-HA:cKO stage V gametocytes treated with DMSO or rapamycin shows a slight but significant reduction in the proportion of male gametocytes upon PfDCX-HA depletion (mean ± SD, n=3 biological replicates, 2-way ANOVA multiple-comparisons correction). **(B)** Representative immunofluorescence image showing two males and one female gametocyte. The female gametocyte is identified by the expression of Pfg377. Scale bar: 10µm.

**Supplementary Tables**

**Table S1: Primers used in this study**

| Primer name | 5' to 3' sequence | observations |
|---|---|---|
| **PfDCX 5' int F** | ACCTCCAGCTGATTTTATGA | Forward primer to check for 5' integration |
| **PfDCX 5' int R** | CGTATAATGTATGCTATACGAACG | Reverse primer to check for 5' integration |
| **PfDCX 3' int F** | GTGCCCGATTATGCGTATCCTTA | Forward primer to check for 3' integration |
| **PfDCX 3' int R** | TCGTTTTGCACTTATCCTCA | Reverse primer to check for 3' integration |
| **PfDCX WT seq F** | GTTGCCTATCCATTCAAACAAATA | Forward primer to check for WT presence |
| **PfDCX WT seq R** | GTTAGAAAGTGAAGACTCAAGTTTC | Reverse primer to check for WT presence |
| **PCR A_F** | ATGGAGAATTTTGATGAAGTT | Forward primer to check for excision of PfDCX-HA |
| **PCR A_R** | CAAAGAAAAATGCATTACTCACAG | Reverse primer to check for excision of PfDCX-HA |
| **PCR B_F** | GCGAGGAGGACAATTTGAAG | Forward primer to check for excision of PfDCX-HA |
| **PCR B_R** | AAGTTACGTATAGGTGCAGGAGG | Reverse primer to check for excision of PfDCX-HA |
| **PfDCX gRNA F5** | ATTGAAGTTTCGAATGGGTGCAGG | Forward primer targeting the endogenous locus |
| **PfDCX gRNA R5** | AAACCCTGCACCCATTCGAAACTT | Reverse primer targeting the endogenous locus |
| **PfDCX gRNA F6** | ATTGGTGAAGACTCAAGTTTCGAA | Forward primer targeting the endogenous locus |
| **PfDCX gRNA R6** | AAACTTCGAAACTTGAGTCTTCAC | Reverse primer targeting the endogenous locus |
| **PbDCX-HA integration F** | TCGTCTGAGTTAGACGGTTC | Forward primer to check for integration |
| **PbDCX-HA integration R** | tggaacatcataagggtaagca | Reverse primer to check for integration |
| **WT presence F** | TTGGGCCCATACGGAAAATA | Forward primer to check for WT presence *in P. berghei* |
| **WT presence R** | CAATCGGGCATGCATCTAAA | Reverse primer to check for WT presence *in P. berghei* |
| **PbDCX-HA gRNA1** | TATTAGATGGTCTATGGATGCTGG | Forward primer targeting the endogenous locus |
| | AAAACCCAGCATCCATAGACCATCT | Reverse primer targeting the endogenous locus |

**Table S2: List of antibodies and dyes**

| Antibody | Fc species | Reference | Source | Dilution | | | |
|---|---|---|---|---|---|---|---|
| | | | | U-ExM | iU-ExM | WB | IFA |
| anti-alpha tubulin | Guinea pig | AA345 | Unige antibody platform | 1/250 | 1/250 | | 1/2000 |
| anti-alpha tubulin | chicken | DM1A | Sigma-Aldrich | | | | 1/500 |
| anti-beta tubulin | Guinea pig | AA344 | Unige antibody platform | 1/250 | 1/250 | | 1/2000 |
| anti-GFP | Rabbit | TP401 | Torrey Pines Biolabs | 1/250 | | | |
| anti-HA | Rat | 3F10 | Roche | 1/250 | 1/250 | 1/1000 | 1/250 |
| anti-Pfs25-cy3 | Mouse | home made | Delves et al., 2013 | | | | 1/1000 |
| anti-actin | Mouse | home made | Herm-Götz et al., 2002 | | | 1/10 | |
| anti-Pfg377 | Rabbit | home made | Alano et al., 1995 | | | | 1/1000 |
| Anti-PolyE | Rabbit | AG-25B-0030 | AdipoGen | 1/400 | | | |
| anti-mouse 488 | Goat | A11001 | ThermoFisher | | | | 1/1000 |
| anti-rat 488 | Goat | A11006 | Invitrogen | 1/400 | 1/400 | | 1/1000 |
| anti-rabbit 568 | Goat | A11011 | Invitrogen | | | | 1/250 |
| anti-rabbit 488 | Goat | A11034 | Invitrogen | 1/400 | | | |
| anti-guinea pig 405 | Goat | Ab175678 | Abcam | 1/400 | | | 1/1000 |
| anti-guinea pig 568 | Goat | A11075 | Invitrogen | | 1/400 | | |
| anti-rat HRP | Goat | 31470 | Invitrogen | | | 1/3000 | |
| anti-mouse HRP | Goat | 32430 | Invitrogen | | | 1/10000 | |

| Dyes | Reference | Source | Final concentration |
|---|---|---|---|
| Atto 594 NHS-ester | 8741 | Merck | 10 µg/mL in PBS 1X for U-ExM |
| Alexa Fluor™ 405 NHS Ester | A30000 | Invitrogen | 2 µg/mL in PBS 1X for iU-ExM |
| SytoxTM Deep Red Nucleic Acid | S11381 | ThermoFisher | 0.5 µM in PBS 1X |
| Hoescht | H3570 | ThermoFisher | 50 µg/mL in PBS 1X |
| mCling | 710 006AT647N | SYSY antibodies | 100nM in PBS 1X |

**Table S3: Chemicals used in U-ExM**

| Designation | Source | Reference |
|---|---|---|
| Formaldehyde 36.5–38% | SIGMA | F8775 |
| Acrylamide (AA, 40%) | SIGMA | A4058 |
| N,N0 -methylenbisacrylamide (BIS, 2%) | SIGMA | M1533 |
| Sodium Acrylate (SA, 97–99%) | SIGMA | 408220 |
| Ammonium persulfate (APS) | ThermoFisher | 17874 |
| Tetramethylethylenediamine (TEMED) | ThermoFisher | 17919 |
| Poly-D-Lysine | Gibco | A3890401 |
| Tween20 BioChemica | AppliChem | A1389,0500 |
| Phosphate Buffered Saline (10x) | SIGMA | D1408-500mL |
| Albumin (BSA) fraction V | AppliChem | A1391,0100 |
| Sodium dodecyl sulfate (SDS) | SIGMA | 71736 |
| TrizmaBase | SIGMA | T1503-1kg |
| Sodium Chloride | SIGMA | 31434-1kg |

**Supplementary Data**

**Data S1. DNA sequences of PfDCX and PbDCX targeting constructs. (A)** Full nucleotide sequence (5' to 3') of the pUC57_PfDCX-HA targeting vector, generated by gene synthesis and experimentally validated by Sanger sequencing. **(B)** Schematic map of the pUC57_PfDCX-HA targeting vector showing the 5' homology region 1 (HR1; 1008 bp), the recodonised PfDCX coding sequence (CSD) with a loxpint module followed by a C-terminal 6xHA tag and a *Lox66* site, and 3' homology region 2 (HR2; 1008 bp). **(C)** Full nucleotide sequence (5' to 3') of the DCX homology-directed repair (HDR) template, PbDCX_PbU6_3HA_gRNA, generated by gene synthesis and experimentally validated by Sanger sequencing. **(D)** Schematic map of the template.

# A

**>pUC57_PfDCX-HA**

TCGCGCGTTTCGGTGATGACGGTGAAAACCTCTGACACATGCAGCTCCCGGAGACGGTCACAGCTTGTCTGTAAGCGGATGCC
GGGAGCAGACAAGCCCGTCAGGGCGCGTCAGCGGGTGTTGGCGGGTGTCGGGGCTGGCTTAACTATGCGGCATCAGAGCAG
ATTGTACTGAGAGTGCACCATATGCGGTGTGAAATACCGCACAGATGCGTAAGGAGAAAATACCGCATCAGGCGCCATTCGC
CATTCAGGCTGCGCAACTGTTGGGAAGGGCGATCGGTGCGGGCCTCTTCGCTATTACGCCAGCTGGCGAAAGGGGGATGTGC
TGCAAGGCGATTAAGTTGGGTAACGCCAGGGTTTTCCCAGTCACGACGTTGTAAAACGACGGCCAGTGAATTCGAGCTCGGTA
CCTCGCGAATGCATCTAGATATTCATATATATTACTTTTAGGTATATTATTATTTATTAATTTTTTTATGTGTCTATTATTTTGTAA
AACATCTCCGATAAGATATATATATATATATATATATATTTATTTATTTATTTATTTATATATATATATTTATTTTTTTGTAAAGTTT
TAACTCTATATTTGAATAATATATCTACATTATTTTAATAATAATTGTATTTAAACGGTAACTGAAAAAATAAAATAAAATAAAA
TAAAACAATGATTTTTTTAAAACGACGTTATGTAAATACTTCTACATAGTTTAGCGGCCGCTAAATCGATATAAATATATATATA

TGAAGATTATGAGGGAATAAAACATTCTCATAAACATATTGAACATATTTAACACATAATAACAAAATAAACGTTTAATATATTT
TATGTCTATGAACACTACCGTTTATCAAAAAACACAATTTAAAAATAAAACGTTCATTATCATAAGTCACAAACCTTTATTTAATT
TTTTAATAAAAATATATTATGAATAGTATATCTTTATATTCAGAGTGATTAAAAATCCTTTTTCTCATGTCAATTATATATATATAT
ATTTTTAAATTACAATGTTTTATAAAAAGATAACTAAGGTTTATTATAAATGTGGTGAACTTATAATATGTTAAGTCTTTTAAAAT
AAAGAAATAATATTAAAAATAAAAAAGATACATATATAAAAAGCCATAAATGTGTTGTCACATTTATTATATTTATAGCACATA
ATTTGATAACTTATATTTTGAAAAAAATAAAAATATAAGTATATATATATATATATATATATATATATTTATTTATTTATTAAGACAA
GCACAAAATGTGCCCATATAAAACACATTCTGTGCAAAAAAGGAAAATGAAAAATATTATACTTAATTACAACTTATAATTACA
TTTGTATAGTATATCTTATTGTATTTAATTATTTTTCACTATATTGTATTGTCAAGTTGTATTATATCAACGTATATATTTTTTAATG
GAGAATTTTGATGAAGTTATAAAAGAATATCAGAAGTATCTAGAGGGAAAAGAAAAGACCCACAAGGTAAATAAAAAAAATA
ATATACATACCGTTCGTATAGCATACATTATACGAAGTTATTATATATGTATATATATATATATTTATATATTTTATATTCTTTTAG
ATAAACAGTTGTAAGCACCCTTGTTGCGAGGAGGACAATTTGAAGTCTGTGAAAAATTATTTCAAGGTTTACCATAAGAAATAT
ATACCATCAATAATACAGAAGAAGAAGAAAGAGAGAAATGAGCATGCACCACCTTTATTGAAGGATATTCAAGACAAGCGTT
GTACTAATGTGTTTGAGAGGTTAAATGATAAGCAGTTTTACACCGGAGTTCAAAAGACCAAATTCATGGAGTTGTTGAAGAAC
AACAAGAATAAGTCAAGTTATTGTTATAATAACATTAAGGTTTTCTCAACTATGCTTAAGAAACCTTGTAACTACGTTGTTACTC
CAGGAACTTTGGGTATACAAAAATATGGTATACAAACTGGACGTCCAAAGACAATATTCTTATTCAATAATGAGAAGAAATAT
GACAAGGGTGTGTACTTCTTGGTGAAGAGTTACATTAAAAATATTAAGAGTCTTTGCTACGAGATTACAAAGATACTTCAACCA
AGTATAGGTCCAACTCGTAAGATTTACGATCAAAACTTCTCTTTGGTTAGGAACGTAAACGATTTGATAAACGGAGGTAAGTAC
TTGTGTACTTCTGGAGATCCTCCTGCACCTATACGTAACTTATCTTTGCACTTTCTTACTTACCCGTACGACGTCCCGGACTACGC
TGGCTATCCCTATGATGTGCCCGATTATGCGTATCCTTACGATGTTCCAGATTATGCCTACCCATACGATGTGCCTGACTATGCC
GGTTACCCTTACGATGTACCAGACTATGCATACCCTTACGATGTACCTGATTACGCCTAGATAACTTCGTATAGCATACATTATA
CGAACGGTATACATAAAATATAAATAAATAAATAAATAAACATATATATATATATTATATATATGTGTAAATGCAATGAGATCA
CCTTTCATAACAAAAAAAAAAAAAAATAAAAAAAAAATAAAATAAGGGTAATGTTTCATATATTGTTGTATATATTTTCCTTTTTTT
ATATTTTTTTTTTTTTTATATATTTTTTTTTTTTTTTATTATTTTTTTTTTTTATATAATATTGTTTCCTTGTATGTGTTCATAATAAAAA
ATAAATCATTTAAAAAATAAAAAAAAAATTAAATAATTAATTATATAAATAAAAATATAAATATAAAAATTTGTACATATATAGG
TATAGGTGGGTAACATCCCAATGTATTTATACATTTAAATAAGTATATATACATATATATTAAAGTTTTTATTCATTTATTTATTTA
TTTTTATTTTTTTTTTTTTATTATTATTGTATTAGATTTTGGCATGTTAGGCAAATATTTCCTCCTTTACAGACAGTACATGCTCGAT
GACTATTGTTTTTTCTTCCCACTCAAATTGGTTCTCGAATGTTTCCCAATTTTTTTTAAGTTGTTCATACTCTTGATAGGATTGGAA
GAGGCGGCCGCTGCAGTAATTTCTTGAAATTTTTGGTTGAATGAATTTTCTACATTTTTTCTTTCTGTGAGTAATGCATTTTTCTT
TGCTGTTTTCATTTTTTCATCTATATTATCGAATTTTTTCGAATAATTATTTAGATTATTTTTAATATCTAGTTTAAGTTTATCATTA
AATAATGACTTTGGATGTGGTCTAAATAAGAATTGAATAAATGATGGTTTTTTAATTCTCATAAGACATTTGCCTTGAAAAGTCC
ATATATGAAATCCTGCTTCACTATTTTCTTCTCTATAACCAGTATTAAAAGCATTTGGATTATTGGAAACAGCTGAGACTAGGAA
TCTTCCACAACGACTCCAGGATATAGAATTAACTAATAGATGTTCATCTTTATGTATAACATCTACTATCGGATCCCGGGCCCGT
CGACTGCAGAGGCCTGCATGCAAGCTTGGCGTAATCATGGTCATAGCTGTTTCCTGTGTGAAATTGTTATCCGCTCACAATTCC
ACACAACATACGAGCCGGAAGCATAAAGTGTAAAGCCTGGGGTGCCTAATGAGTGAGCTAACTCACATTAATTGCGTTGCGCT
CACTGCCCGCTTTCCAGTCGGGAAACCTGTCGTGCCAGCTGCATTAATGAATCGGCCAACGCGCGGGGAGAGGCGGTTTGCGT
ATTGGGCGCTCTTCCGCTTCCTCGCTCACTGACTCGCTGCGCTCGGTCGTTCGGCTGCGGCGAGCGGTATCAGCTCACTCAAAG
GCGGTAATACGGTTATCCACAGAATCAGGGGATAACGCAGGAAAGAACATGTGAGCAAAAGGCCAGCAAAAGGCCAGGAAC
CGTAAAAAGGCCGCGTTGCTGGCGTTTTTCCATAGGCTCCGCCCCCCTGACGAGCATCACAAAAATCGACGCTCAAGTCAGAG
GTGGCGAAACCCGACAGGACTATAAAGATACCAGGCGTTTCCCCCTGGAAGCTCCCTCGTGCGCTCTCCTGTTCCGACCCTGCC
GCTTACCGGATACCTGTCCGCCTTTCTCCCTTCGGGAAGCGTGGCGCTTTCTCATAGCTCACGCTGTAGGTATCTCAGTTCGGTG
TAGGTCGTTCGCTCCAAGCTGGGCTGTGTGCACGAACCCCCCGTTCAGCCCGACCGCTGCGCCTTATCCGGTAACTATCGTCTT
GAGTCCAACCCGGTAAGACACGACTTATCGCCACTGGCAGCAGCCACTGGTAACAGGATTAGCAGAGCGAGGTATGTAGGCG
GTGCTACAGAGTTCTTGAAGTGGTGGCCTAACTACGGCTACACTAGAAGAACAGTATTTGGTATCTGCGCTCTGCTGAAGCCA
GTTACCTTCGGAAAAAGAGTTGGTAGCTCTTGATCCGGCAAACAAACCACCGCTGGTAGCGGTGGTTTTTTTGTTTGCAAGCA
GCAGATTACGCGCAGAAAAAAAGGATCTCAAGAAGATCCTTTGATCTTTTCTACGGGGTCTGACGCTCAGTGGAACGAAAACT
CACGTTAAGGGATTTTGGTCATGAGATTATCAAAAAGGATCTTCACCTAGATCCTTTTAAATTAAAAATGAAGTTTTAAATCAAT
CTAAAGTATATATGAGTAAACTTGGTCTGACAGTTACCAATGCTTAATCAGTGAGGCACCTATCTCAGCGATCTGTCTATTTCGT
TCATCCATAGTTGCCTGACTCCCCGTCGTGTAGATAACTACGATACGGGAGGGCTTACCATCTGGCCCCAGTGCTGCAATGATA
CCGCGAGACCCACGCTCACCGGCTCCAGATTTATCAGCAATAAACCAGCCAGCCGGAAGGGCCGAGCGCAGAAGTGGTCCTG

CAACTTTATCCGCCTCCATCCAGTCTATTAATTGTTGCCGGGAAGCTAGAGTAAGTAGTTCGCCAGTTAATAGTTTGCGCAACG
TTGTTGCCATTGCTACAGGCATCGTGGTGTCACGCTCGTCGTTTGGTATGGCTTCATTCAGCTCCGGTTCCCAACGATCAAGGC
GAGTTACATGATCCCCCATGTTGTGCAAAAAAGCGGTTAGCTCCTTCGGTCCTCCGATCGTTGTCAGAAGTAAGTTGGCCGCAG
TGTTATCACTCATGGTTATGGCAGCACTGCATAATTCTCTTACTGTCATGCCATCCGTAAGATGCTTTTCTGTGACTGGTGAGTA
CTCAACCAAGTCATTCTGAGAATAGTGTATGCGGCGACCGAGTTGCTCTTGCCCGGCGTCAATACGGGATAATACCGCGCCAC
ATAGCAGAACTTTAAAAGTGCTCATCATTGGAAAACGTTCTTCGGGGCGAAAACTCTCAAGGATCTTACCGCTGTTGAGATCCA
GTTCGATGTAACCCACTCGTGCACCCAACTGATCTTCAGCATCTTTTACTTTCACCAGCGTTTCTGGGTGAGCAAAAACAGGAA
GGCAAAATGCCGCAAAAAAGGGAATAAGGGCGACACGGAAATGTTGAATACTCATACTCTTCCTTTTTCAATATTATTGAAGC
ATTTATCAGGGTTATTGTCTCATGAGCGGATACATATTTGAATGTATTTAGAAAAATAAACAAATAGGGGTTCCGCGCACATTT
CCCCGAAAAGTGCCACCTGACGTCTAAGAAACCATTATTATCATGACATTAACCTATAAAAATAGGCGTATCACGAGGCCCTTT
CGTC

**B**

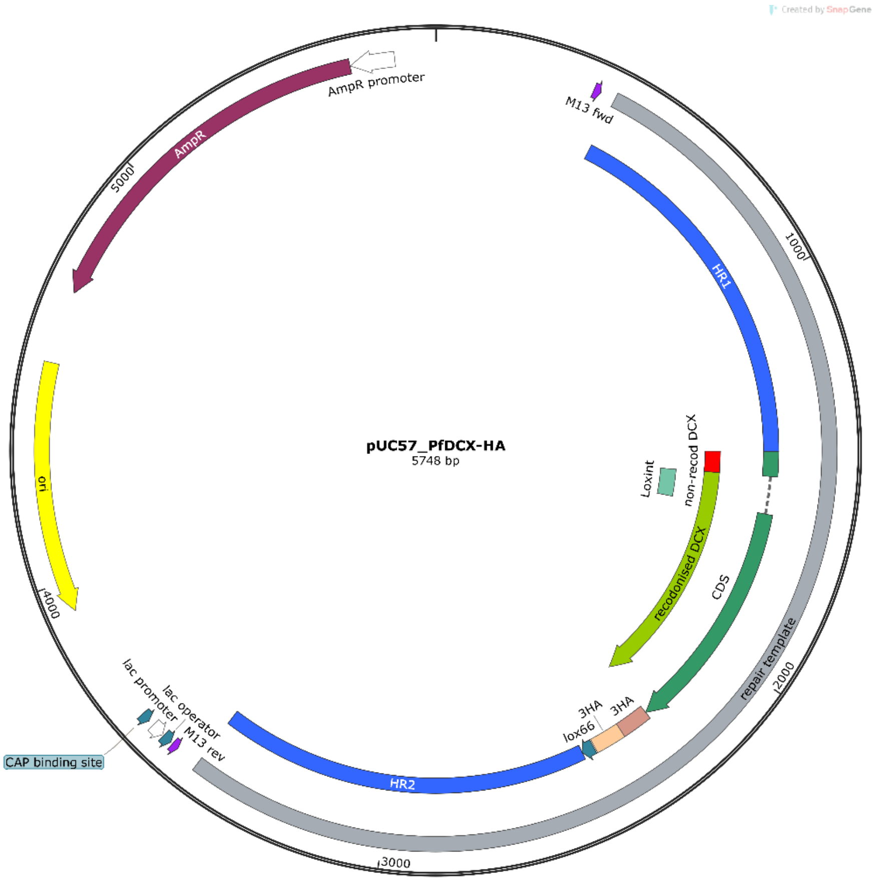


**C**

**>PbDCX homology-directed repair template (PbDCX_PbU6_3HA_gRNA)**

ttgagatcctttttttctgcgcgtaatctgctgcttgcaaacaaaaaaaccaccgctaccagcggtggtttgtttgccggatcaagagctaccaactcttt
ttccgaaggtaactggcttcagcagagcgcagataccaaatactgtccttctagtgtagccgtagttaggccaccacttcaagaactctgtagcaccgc
ctacatacctcgctctgctaatcctgttaccagtggctgctgccagtggcgataagtcgtgtcttaccgggttggactcaagacgatagttaccggataa
ggcgcagcggtcgggctgaacggggggttcgtgcacacagcccagcttggagcgaacgacctacaccgaactgagatacctacagcgtgagcattg
agaaagcgccacgcttcccgaagggagaaaggcggacaggtatccggtaagcggcagggtcggaacaggagagcgcacgagggagcttccaggg
ggaaacgcctggtatctttatagtcctgtcgggtttcgccacctctgacttgagcgtcgatttttgtgatgctcgtcaggggggcggagcctatggaaaa
acgccagcaacgcggcctttttacggttcctggccttttgctggccttttgctcacatgttctttcctgcgttatcccctgattctgtggataaccgtattacc
gcctttgagtgagctgataccgctcgccgcagccgaacgaccgagcgcagcgagtcagtgagcgaggaagcggaagagcgcccaatacgcaaacc

gcctctccccgcgcgttggccgattcattaatgcagctggcacgacaggtttcccgactggaaagcgggcagtgagcgcaacgcaattaatgtgagtt
agctcactcattaggcaccccaggctttacactttatgcttccggctcgtatgttgtgtggaattgtgagcggataacaatttcacacaggaaacagcta
tgaccatgattacgccaagcttgaaaaaaaaagaaacaaacaatataaaagtgtatttgaacgtttaacagaccaaaacttttatacaggaactcat
aaaaaaaaattcgaggcattggtaaaatgtaaattgaatgaacaggatatggataattatttgtctataaatttagaaatgcatacaaatataaaaaa
tgaaaacatgatttctcaatattctttttcttctattggaataaatcaattcgaaaatggaaagaaaaaaaaaaaaaaagaaaaaaatttagtagttac
cccaggaatacttggtatacaaaaatatggaattcaaattgctcgccccaaaagtatttggctatatcgaaatggagataaacatcataatggcttact
tttctttattaagtcccatataaataattttaaattgttattgttcgaaattacaaaagtattaaatcctatcattgggcccatacggaaaatatatgatca
aaattttagactcataaaaaatatacaacaactcaacgatggttccaaatatttatgcacctctggtgatccacctgcttcaattgatcatttaggaaag
ttcaaatccaagtgggttatacaagggtatccttatgatgtaccagattatgcatatccatacgatgtacctgattatgcttacccttatgatgttccaga
ctacgcatagcatattatagtttaaggaaaagtagtgaaattgtcaaaatatcccacaaaatcatttagaaagagatgataaataaatatgaactcctt
tttagatgcatgcccgattgtccatatattttgcaacaaaatgaagtgtatatggtatttcccaatctgaaccatattctacattccttggggaacacact
gatatttattttaaaaattaaaaatatttcttgtattaaagttttttgttatgtgcgtagtagtttacattagcgtcattacgtgtaaagaaatataaaaagt
taaaaaaaaaataaaaaataaaaaaggacactaataaatgatgcaaaaaaagagaagcagcatagaaacttctcaaaactcagtaaaatgtacat
catttctatccaaacatacacaaatataaatgaatatataagagaaaatctatgcgaagatttcttgtttaacagaaaggacatgttcaataacaactg
tttttcttcccactcaaattggctttcaaatgtttcccaatttctcctgaacaattcatactctttaaaggattggaataatcttgtgattttctccaatttt
cattaaatccatggtccccgcggacgctaatcgtagctagcctgctgactcgaggaattccttaagtttacaatttaatattcatactttaagtatttttg
tagtatcctagatattgtgctttaaatgctcacccctcaaagcaccagtaatattttcatccactgaaataccattaaattttcaaaaaaatactatgcat
ataatgttatacatataaacataaaacgccatgtaaatcaaaaaatatataaaaatatgtataaaaataaatatgcactaaatataagctaattatgc
ataaaaattaaagtgccctttattaactagtcgtaattatttatatttctatgttataaaaaaatcctcatataataatataattaatatatgtaatgttttt
tttattttataattttaatataaaataatatgtaaattaattcaaaaaataaatataattgttgtgaaacaaaaaacgtaatttttcatttgccttcaaaa
tttaaatttattttaatatttcctaaaatatatatactttgtgtataaatatataaaaatatatatttgcttataaataaataaaaaattttataaaacatag
ggggatctatggttggttcgctaaactgcatcgtcgctgtgtcccagaacatgggcatcggcaagaacggggacctgccctggccaccgctcaggaac
gaatttagatatttccagagaatgaccacaacctcttcagtagaaggtaaacagaatctggtgattatgggtaagaagacctggttctccattcctgag
aagaatcgacctttaaagggtagaattaatttagttctcagcagagaactcaaggaacctccacaaggagctcattttctttccagaagtctagatgat
gccttaaaacttactgaacaaccagaattagcaaataaagtagacatggtctggatagttggtggcagttctgtttataaggaagccatgaatcaccc
aggccatcttaaactatttgtgacaaggatcatgcaagactttgaaagtgacacgtttttttccagaaattgatttggagaaatataaacttctgccagaa
tacccaggtgttctctctgatgtccaggaggagaaaggcattaagtacaaatttgaagtatatgagaagaatgatgcaagcggaggaggtggatctg
gtggaggtggaagtgcaagcgtgacagggggaatggctagcaagtgggatcagaagggtatggacattgcctatgaggaggcggccttaggttaca
aagagggtggtgttcctattggcggatgtcttatcaataacaaagacggaagtgttctcggtcgtggtcacaacatgagatttcaaaagggatccgcc
acactacatggtgagatctccactttggaaaactgtgggagattagagggcaaagtgtacaaagataccactttgtatacgacgctgtctccatgcga
catgtgtacaggtgccatcatcatgtatggtattccacgctgtgttgtcggtgagaacgttaatttcaaaagtaagggcgagaaatatttacaaactag
aggtcacgaggttgttgttgttgacgatgagaggtgtaaaaagatcatgaaacaatttatcgatgaaagacctcaggattggtttgaagatattggtga
ggcttcggaaccatttaagaacgtctacttgctacctcaaacaaaccaattgctgggtttgtacaccatcatcagaaataagaatacaactagacctga
tttcattttctactccgatagaatcatcagattgttggttgaagaaggtttgaaccatctacctgtgcaaaagcaaattgtggaaactgacaccaacgaa
aacttcgaaggtgtctcattcatgggtaaaatctgtggtgtttccattgtcagagctggtgaatcgatggagcaaggattaagagactgttgtaggtctg
tgcgtatcggtaaaattttaattcaaagggacgaggagactgctttaccaaagttattctacgaaaaattaccagaggatatatctgaaaggtatgtct
tcctattagacccaatgctggccaccggtggtagtgctatcatggctacagaagtcttgattaagagaggtgttaagccagagagaatttacttcttaa
acctaatctgtagtaaggaagggattgaaaaataccatgccgccttcccagaggtcagaattgttactggtgccctcgacagaggtctagatgaaaac
aagtatctagttccagggttgggtgactttggtgacagatactactgtgttggttcgggagagggcagaggatccctgctaacatgcggtgatgtcgag
gagaatcctggcccagaatcgatggactataaggaccacgacggagactacaaggatcatgatattgattacaaagacgatgacgataagatggcc
ccaaagaagaagcggaaggtcggtatccacggagtcccagcagccgacaagaagtacagcatcggcctggacatcggcaccaactctgtgggctgg
gccgtgatcaccgacgagtacaaggtgcccagcaagaaattcaaggtgctgggcaacaccgaccggcacagcatcaagaagaacctgatcggagc
cctgctgttcgacagcggcgaaacagccgaggccacccggctgaagagaaccgccagaagaagatacaccagacggaagaaccggatctgctatc
tgcaagagatcttcagcaacgagatggccaaggtggacgacagcttcttccacagactggaagagtccttcctggtggaagaggataagaagcacg

agcggcaccccatcttcggcaacatcgtggacgaggtggcctaccacgagaagtaccccaccatctaccacctgagaaagaaactggtggacagca
ccgacaaggccgacctgcggctgatctatctggccctggcccacatgatcaagttccggggccacttcctgatcgagggcgacctgaaccccgacaac
agcgacgtggacaagctgttcatccagctggtgcagacctacaaccagctgttcgaggaaaaccccatcaacgccagcggcgtggacgccaaggcc
atcctgtctgccagactgagcaagagcagacggctggaaaatctgatcgcccagctgcccggcgagaagaagaatggcctgttcggaaacctgattg
ccctgagcctgggcctgacccccaacttcaagagcaacttcgacctggccgaggatgccaaactgcagctgagcaaggacacctacgacgacgacct
ggacaacctgctggcccagatcggcgaccagtacgccgacctgtttctggccgccaagaacctgtccgacgccatcctgctgagcgacatcctgagag
tgaacaccgagatcaccaaggcccccctgagcgcctctatgatcaagagatacgacgagcaccaccaggacctgaccctgctgaaagctctcgtgcg
gcagcagctgcctgagaagtacaaagagattttcttcgaccagagcaagaacggctacgccggctacattgacggcggagccagccaggaagagtt
ctacaagttcatcaagcccatcctggaaaagatggacggcaccgaggaactgctcgtgaagctgaacagagaggacctgctgcggaagcagcggac
cttcgacaacggcagcatccccccaccagatccacctgggagagctgcacgccattctgcggcggcaggaagatttttacccattcctgaaggacaacc
gggaaaagatcgagaagatcctgaccttccgcatcccctactacgtgggccctctggccaggggaaacagcagattcgcctggatgaccagaaaga
gcgaggaaaccatcaccccctggaacttcgaggaagtggtggacaagggcgcttccgcccagagcttcatcgagcggatgaccaacttcgataaga
acctgcccaacgagaaggtgctgcccaagcacagcctgctgtacgagtacttcaccgtgtataacgagctgaccaaagtgaaatacgtgaccgaggg
aatgagaaagcccgccttcctgagcggcgagcagaaaaaggccatcgtggacctgctgttcaagaccaaccggaaagtgaccgtgaagcagctgaa
agaggactacttcaagaaaatcgagtgcttcgactccgtggaaatctccggcgtggaagatcggttcaacgcctccctgggcacataccacgatctgc
tgaaaattatcaaggacaaggacttcctggacaatgaggaaaacgaggacattctggaagatatcgtgctgaccctgacactgtttgaggacagaga
gatgatcgaggaacggctgaaaacctatgcccacctgttcgacgacaaagtgatgaagcagctgaagcggcggagatacaccggctggggcaggct
gagccggaagctgatcaacggcatccgggacaagcagtccggcaagacaatcctggatttcctgaagtccgacggcttcgccaacagaaacttcatg
cagctgatccacgacgacagcctgacctttaaagaggacatccagaaagcccaggtgtccggccagggcgatagcctgcacgagcacattgccaatc
tggccggcagccccgccattaagaagggcatcctgcagacagtgaaggtggtggacgagctcgtgaaagtgatgggccggcacaagcccgagaac
atcgtgatcgaaatggccagagagaaccagaccacccagaagggacagaagaacagccgcgagagaatgaagcggatcgaagagggcatcaaa
gagctgggcagccagatcctgaaagaacaccccgtggaaaacacccagctgcagaacgagaagctgtacctgtactacctgcagaatggcgggat
atgtacgtggaccaggaactggacatcaaccggctgtccgactacgatgtggaccatatcgtgcctcagagctttctgaaggacgactccatcgacaa
caaggtgctgaccagaagcgacaagaaccggggcaagagcgacaacgtgccctccgaagaggtcgtgaagaagatgaagaactactggcggcag
ctgctgaacgccaagctgattacccagagaaagttcgacaatctgaccaaggccgagagaggcggcctgagcgaactggataaggccggcttcatc
aagagacagctggtggaaacccggcagatcacaaagcacgtggcacagatcctggactcccggatgaacactaagtacgacgagaatgacaagct
gatccgggaagtgaaagtgatcaccctgaagtccaagctggtgtccgatttccggaaggatttccagttttacaaagtgcgcgagatcaacaactacc
accacgcccacgacgcctacctgaacgccgtcgtgggaaccgccctgatcaaaaagtaccctaagctggaaagcgagttcgtgtacggcgactacaa
ggtgtacgacgtgcggaagatgatcgccaagagcgagcaggaaatcggcaaggctaccgccaagtacttcttctacagcaacatcatgaactttttca
agaccgagattaccctggccaacggcgagatccggaagcggcctctgatcgagacaaacggcgaaaccggggagatcgtgtgggataagggccgg
gattttgccaccgtgcggaaagtgctgagcatgccccaagtgaatatcgtgaaaaagaccgaggtgcagacaggcggcttcagcaaagagtctatcc
tgcccaagaggaacagcgataagctgatcgccagaaagaaggactgggaccctaagaagtacggcggcttcgacagccccaccgtggcctattctg
tgctggtggtggccaaagtggaaaagggcaagtccaagaaactgaagagtgtgaaagagctgctggggatcaccatcatggaaagaagcagcttcg
agaagaatcccatcgactttctggaagccaagggctacaaagaagtgaaaaaggacctgatcatcaagctgcctaagtactccctgttcgagctgga
aaacggccggaagagaatgctggcctctgccggcgaactgcagaagggaaacgaactggccctgccctccaaatatgtgaacttcctgtacctggcc
agccactatgagaagctgaagggctcccccgaggataatgagcagaaacagctgtttgtggaacagcacaagcactacctggacgagatcatcgag
cagatcagcgagttctccaagagagtgatcctggccgacgctaatctggacaaagtgctgtccgcctacaacaagcaccgggataagcccatcagag
agcaggccgagaatatcatccacctgtttaccctgaccaatctgggagcccctgccgccttcaagtactttgacaccaccatcgaccggaagaggtac
accagcaccaaagaggtgctggacgccaccctgatccaccagagcatcaccggcctgtacgagacacggatcgacctgtctcagctgggaggcgac
aaaaggccggcggccacgaaaaaggccggccaggcaaaaaagaaaaagtaagaattaccggtgatcccgtttttcttacttatatatttataccaatt
gattgtatttataactgtaaaaatgtgtatgttgtgtgcatatttttttttgtgcatgcacatgcatgtaaatagctaaaattatgaacattttattttttgtt
cagaaaaaaaaaactttacacacataaaatggctagtatgaatagccatattttatataaattaaatcctatgaatttatgaccatattaaaaatttag
atatttatggaacataatatgtttgaaacaataagacaaaattattattattattatttttactgttataattatgttgtctcttcaatgattcataaata
gttggacttgattttaaaatgtttataatatgattagcatagttaaataaaaaaagttgaaaaattaaaaaaaaacatataaacacaaatgatgttttt

tccttcaatttcggcactggccgtcgttttacaacgtcgtgactgggaaaaccctggcgttacccaacttaatcgccttgcagcacatccccctttcgcca
gctggcgtaatagcgaagaggcccgcaccgatcgcccttcccaacagttgcgcagcctgaatggcgaatgatcgagaactattgtttcctttttgttttt
ttatttttaattaaaatgtaaatataaactcacaaaatgaatgaataaataaataaataaataaataaaaaatgttaaatcgtgttaaaccgcgtgctt
catgtgatatggcaagcgattaatggccattttatccatttttataatttgataaaaaattaaaacaaataaatgtaatgttaattatattttgtactataa
atattattctctacttcacgcaaatgtaacactatccaactacactcataatgtttaaagaaaattggcattgttttaccatttttttaaaacaaaaaaag
aataaaatactttcgtattatttagctaaaattgtactaaatttcataccaaaaaaaaatatatatatgtatgcagctaatatttatttgataagtttttat
gtaaataatatttaattaaaaaaaaaaattacatagcatgccgaatgcactattcattttatgggggtaatttttttgagtaaccatttttactttttttttaca
tatgcgcatacttcgagttatacaatattattagatggtctatggatgctgggttttagagctagaaatagcaagttaaaataaggctagtccgttatca
acttgaaaaagtggcaccgagtcggtgctttttttggacgtcaggtggcacttttcggggaaatgtgcgcggaacccctatttgtttatttttctaaatac
attcaaatatgtatccgctcatgagacaataaccctgataaatgcttcaataatattgaaaaaggaagagtatgagtattcaacatttccgtgtcgccct
tattcccttttttgcggcattttgccttcctgtttttgctcacccagaaacgctggtgaaagtaaaagatgctgaagatcagttgggtgcacgagtgggtt
acatcgaactggatctcaacagcggtaagatccttgagagttttcgccccgaagaacgttttccaatgatgagcacttttaaagttctgctatgtggcgc
ggtattatcccgtattgacgccgggcaagagcaactcggtcgccgcatacactattctcagaatgacttggttgagtactcaccagtcacagaaaagc
atcttacggatggcatgacagtaagagaattatgcagtgctgccataaccatgagtgataacactgcggccaacttacttctgacaacgatcggagga
ccgaaggagctaaccgcttttttgcacaacatgggggatcatgtaactcgccttgatcgttgggaaccggagctgaatgaagccataccaaacgacga
gcgtgacaccacgatgcctgtagcaatggcaacaacgttgcgcaaactattaactggcgaactacttactctagcttcccggcaacaattaatagact
ggatggaggcggataaagttgcaggaccacttctgcgctcggcccttccggctggctggtttattgctgataaatctggagccggtgagcgtgggtctc
gcggtatcattgcagcactggggccagatggtaagccctcccgtatcgtagttatctacacgacggggagtcaggcaactatggatgaacgaaatag
acagatcgctgagataggtgcctcactgattaagcattggtaactgtcagaccaagtttactcatatatactttagattgatttaaaacttcatttttaatt
taaaaggatctaggtgaagatcctttttgataatctcatgaccaaaatcccttaacgtgagttttcgttccactgagcgtcagaccccgtagaaaagatc
aaaggatcttc

**D**

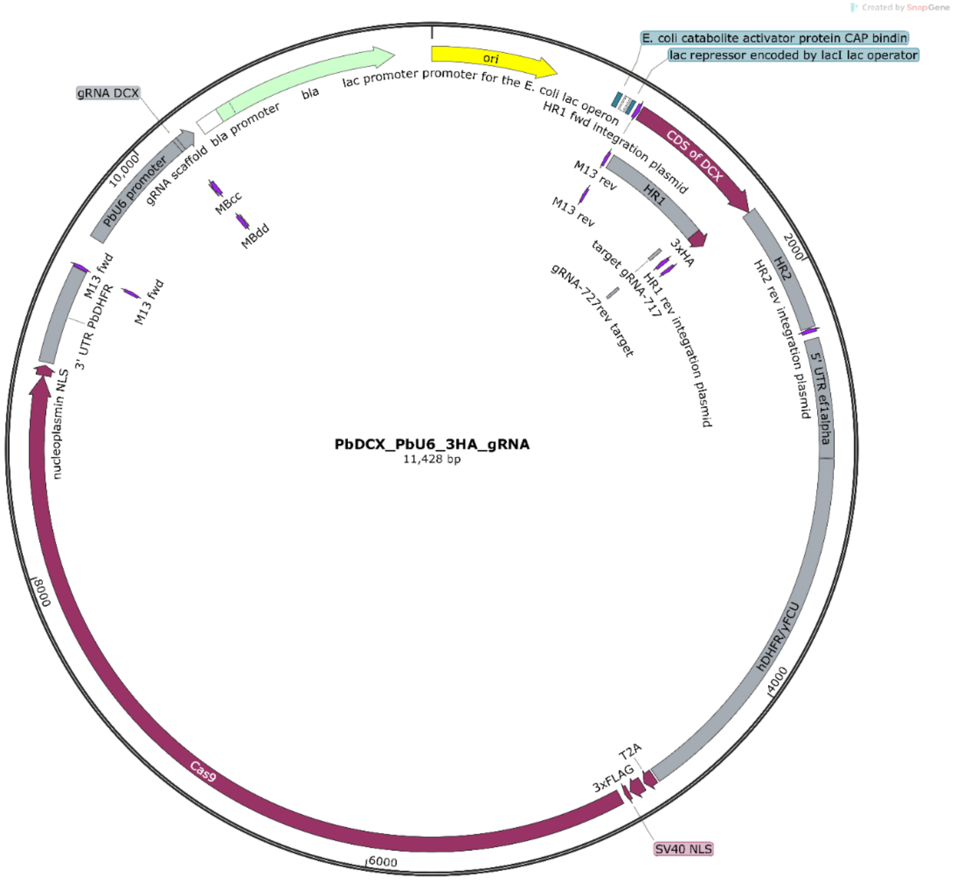

PbDCX_PbU6_3HA_gRNA
11,428 bp
ori
E. coli catabolite activator protein CAP bindin
lac repressor encoded by lacI lac operator
lac promoter promoter for the E. coli lac operon
HR1 fwd integration plasmid
CDS of DCX
M13 rev
M13 rev
HR1
3xHA
target gRNA-717
gRNA-727rev target
HR1 rev integration plasmid
HR2
HR2 rev integration plasmid
2000
5' UTR ef1alpha
hDHFR/yFCU
4000
T2A
3xFLAG
SV40 NLS
6000
Cas9
8000
nucleoplasmin NLS
3' UTR PbDHFR
M13 fwd
M13 fwd
PbU6 promoter
10,000
gRNA DCX
gRNA scaffold
bla promoter
bla
MBcc
MBdd
Created by SnapGene

**Data S2: Uncropped Western blot for Fig. 1A**

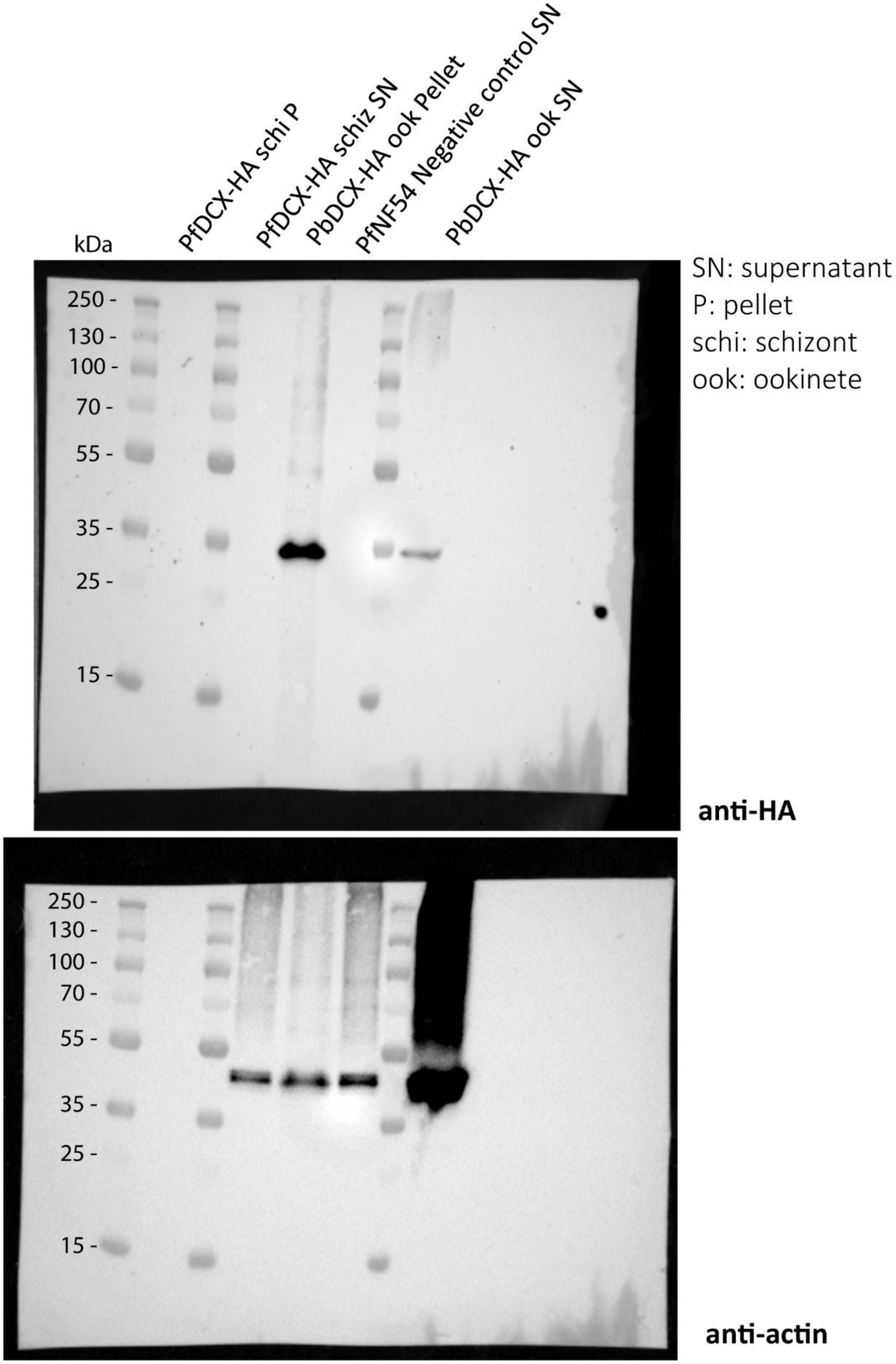

**Data S3: Uncropped Western blot for Fig. 2B**

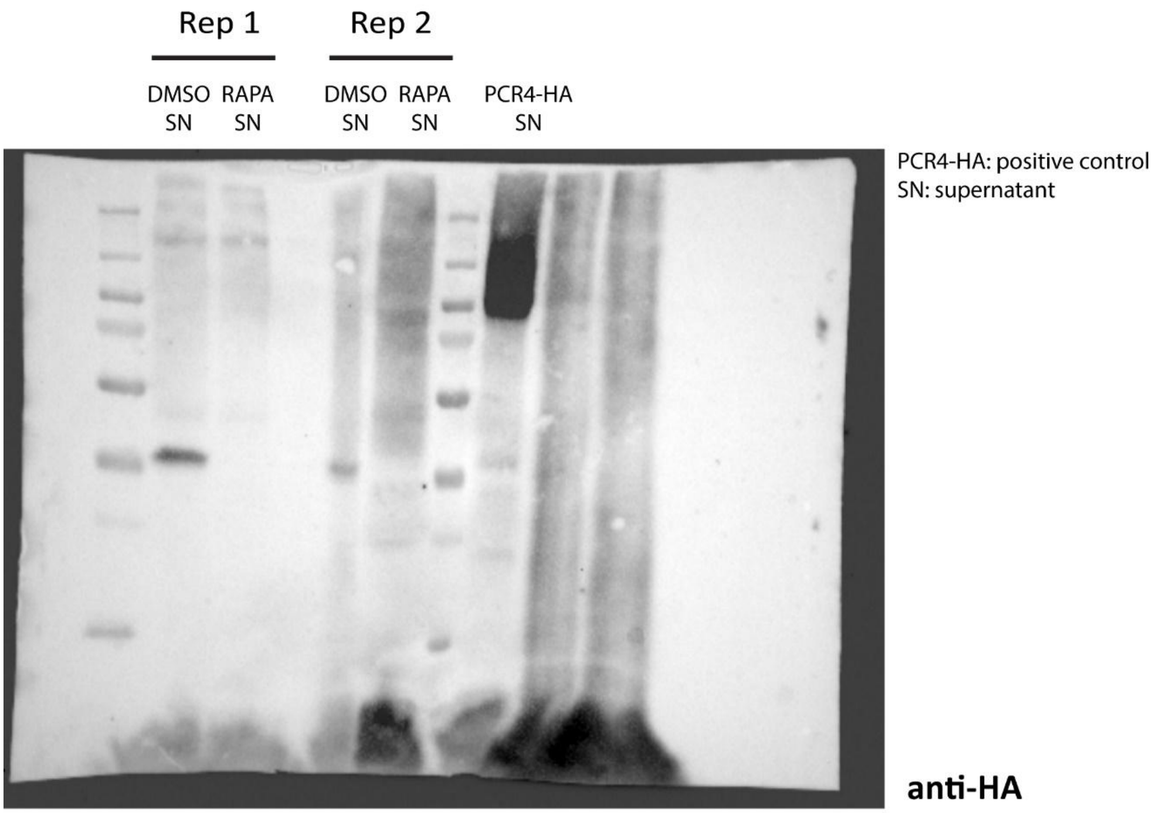


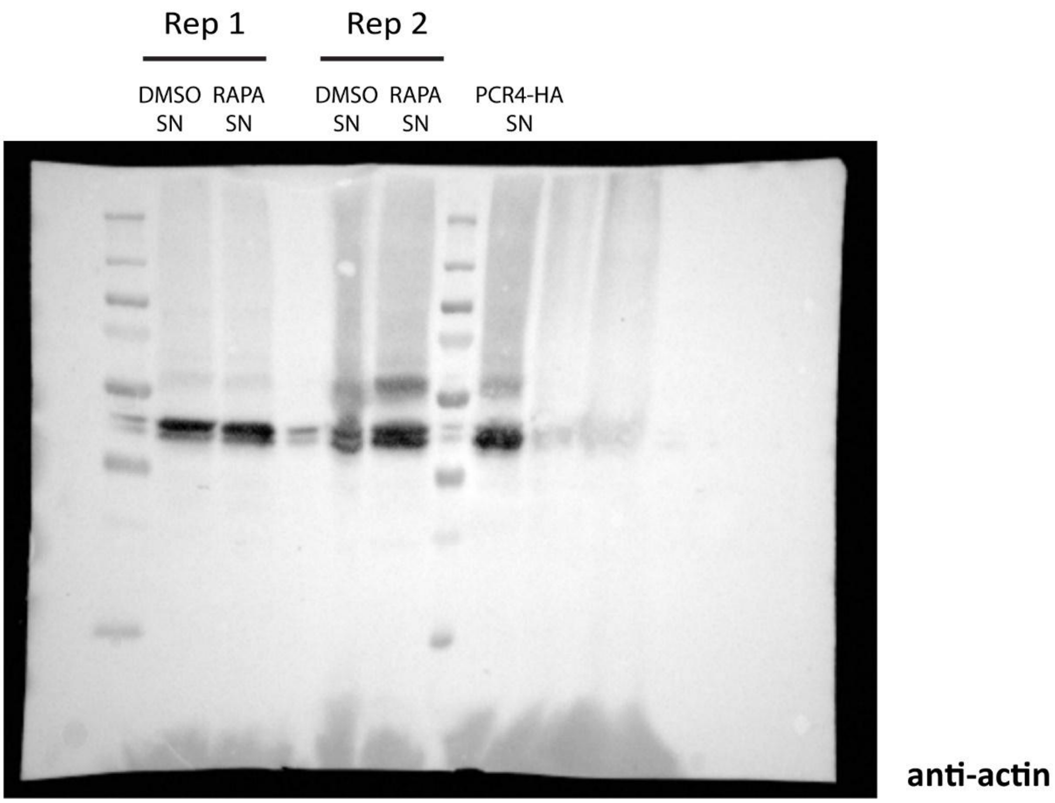
